# Predicting MEG brain functional connectivity using microstructural information

**DOI:** 10.1101/2020.09.15.298307

**Authors:** Eirini Messaritaki, Sonya Foley, Simona Schiavi, Lorenzo Magazzini, Bethany Routley, Derek K. Jones, Krish D. Singh

**Affiliations:** Cardiff University Brain Research Imaging Centre (CUBRIC), School of Psychology, Cardiff University, Maindy Road, Cardiff, CF24 4HQ, UK; BRAIN Biomedical Research Unit, Cardiff University, Maindy Road, Cardiff, CF24 4HQ, UK; Division of Psychological Medicine and Clinical Neurosciences, School of Medicine, Cardiff University, Cardiff, CF24 4HQ, UK; School of Psychology, Cardiff University, 70 Park Place, Cardiff, CF10 3AT, UK; Department of Computer Science, University of Verona, Verona, Italy

## Abstract

Understanding how human brain microstructure influences functional connectivity is an important endeavor. In this work, magnetic resonance imaging data from ninety healthy participants were used to calculate structural connectivity matrices using the streamline count, fractional anisotropy, radial diffusivity and a myelin measure (derived from multicomponent relaxometry) to assign connection strength. Unweighted binarized structural connectivity matrices were also constructed. Magnetoencephalography resting-state data from those participants were used to calculate functional connectivity matrices, via correlations of the Hilbert envelopes of beamformer timeseries at the delta, theta, alpha and beta frequency bands. Non-negative matrix factorization was performed to identify the components of the functional connectivity. Shortest-path-length and search-information analyses of the structural connectomes were used to predict functional connectivity patterns for each participant.

The microstructure-informed algorithms predicted the components of the functional connectivity more accurately than they predicted the total functional connectivity. This provides a methodology for better understanding of functional mechanisms. The shortest-path-length algorithm exhibited the highest prediction accuracy. Of the weights of the structural connectivity matrices, the streamline count and the myelin measure gave the most accurate predictions, while the fractional anisotropy performed poorly. Overall, different structural metrics paint very different pictures of the structural connectome and its relationship to functional connectivity.

## 1 Introduction

The representation of the human brain as a network, in which cortical and subcortical areas (nodes) communicate via white matter tracts that carry neuronal signals (connections or edges), has been used extensively to study the brains of healthy people and of patients that suffer from neurological and neuropsychiatric conditions (for example Hagmann et al. (2008); Griffa et al. (2013); Caeyenberghs and Leemans (2014); Fischer et al. (2014); van den Heuvel and Fornito (2014); Yuan et al. (2014); Baker et al. (2015); Collin et al. (2016); Drakesmith, Caeyenberghs, Dutt, Zammit, Evans, Reichenberg, Lewis, David and Jones (2015); Aerts et al. (2016); Nelson et al. (2017); Vidaurre et al. (2018); Imms et al. (2019)). Structural networks can be derived from diffusion magnetic resonance imaging (MRI) data via tractography methods (Basser et al., 2000; Mukherjee et al., 2008b,a), and represent the intricate wiring of the human brain that allows communication between different brain areas. Functional networks can be constructed from magnetoencephalography (MEG), electroencephalography (EEG) or functional MRI (fMRI) data, by calculating correlations of brain activity between different brain areas (Biswal et al., 1997; Greicius et al., 2003; Brookes et al., 2011), and represent the possible association of physiological activity in those areas. Comparing structural and functional networks can lead to an understanding of the role of the structural connectome on the evocation of functional connectivity, both in healthy and in diseased brains. It can also shed light into whether, in diseased brains, the local synaptic disruptions and the excitation-inhibition imbalance, and the resulting disrupted functional connectome, lead to structural impairments, or whether it is the structural impairments that lead to functional deficiencies. Such knowledge can inform possible interventions that target structural or functional deficiencies in patients (Friston et al., 2016).

The effort to relate the human structural and functional connectomes started several years ago. Honey et al. (2009) investigated whether systems-level properties of fMRI-derived functional networks can be accounted for by properties of structural networks, and found that although resting-state functional connectivity (i.e. connectivity in the absence of a task) is frequently present between regions without direct structural connections, its strength, persistence, and spatial patterns are constrained by the large-scale anatomical structure of the human brain. Gõni et al. (2014) used analytic measures of network communication on the structural con-nectome of the human brain and explored the capacity of these measures to predict restingstate functional connectivity derived from functional MRI data. Shen et al. (2015) showed that resting-state functional connectivity (measured using fMRI) is partly dependent on direct structural connections, but also that dynamic coordination of activity can occur via polysynaptic pathways, as initially postulated by Robinson (2012). Mišić et al. (2016) used singular value decomposition to calculate covariance between the structural and functional (measured with fMRI) connections in data from the Human Connectome Project (Essen et al., 2013), concluding that functional connectivity patterns do not conform to structural connectivity patterns, and that network-wide interactions involve structural connections that do not exist within the functional network in question. Mill et al. (2017) argued that it is essential to consider the temporal variability of resting-state functional connectivity, because that leads to a better understanding of the components that represent task-based connectivity, and therefore of how task-based connectivity can emerge from the structural connectome. Tewarie et al. (2020) related the approach in which functional connectivity is explained by all possible paths in the structural network, i.e. series expansion approach, to the eigenmode approach. There is also a growing literature on forward generative mechanisms that link microstructure to function (for example Honey et al. (2007, 2009); Deco et al. (2011, 2014, 2017)). Finally, Cabral et al. (2017) give a comprehensive review on the subject, while Suarez et al. (2020) argue that structural connectomes that are enriched with biological details such as local molecular and cellular data have a better chance of disentangling the relationship between structure and function in the human brain.

Linking functional and structural connectivity has also been done for electrophysiologically measured functional connectivity. Cabral et al. (2014) investigated the mechanisms of MEG resting-state functional connectivity using a model of coupled oscillators on structural brain networks. Garcés et al. (2016) compared networks derived via diffusion-weighted imaging, fMRI and MEG, for nine participants, observing some similarities and some differences in the hub-ness of the nodes and the patterns of functional connectivity. Tewarie et al. (2014) used the structural degree and the Euclidean distance to predict functional networks derived from fMRI and MEG recordings, using a structural network derived from a cohort that was independent of the one for which the functional recordings had been obtained. Pineda-Pardo et al. (2014) described an estimation of functional connectivity derived via MEG recordings, by structural connectivity, and showed that the methodology can serve to classify participants with mild cognitive impairment. Meier et al. (2016) used group-averaged connectomes derived from MEG, fMRI and diffusion-weighted imaging, to investigate the structure-function mapping. Tewarie et al. (2019) proposed a decomposition of the structural connectome into eigenmodes and used those to predict MEG-derived functional connectivity in different frequency bands.

In this work, we are interested in understanding how the brain microstructure relates to electrophysiological functional connectivity in the human brain. We used MRI and MEG data from ninety healthy participants, a much larger sample than used in most previous studies. Specifically, we used diffusion MRI data to construct their white matter tracts and to derive their structural connectivity networks. Although diffusion MRI is a good method of identifying those tracts, it is insensitive to their myelin content. Given that myelin is believed to play a major role in the formation of functional connectivity in the brain, we also used mcDESPOT data (Deoni et al., 2008) to derive a myelin measure to assign strength to the connections of the structural networks. We used the MEG resting-state recordings to calculate correlations between the electrophysiological activity in different brain areas and derive the resting-state functional connectivity networks of the same participants. We then used the structural networks to predict patterns of functional connectivity using algorithms proposed by Gõni et al. (2014). These algorithms have been used successfully to predict fMRI resting-state connectivity in the past and have the benefit that they do not require any parameter fitting, unlike other methods. We postulated that the simple dynamics involved in those algorithms can be effective at capturing MEG resting-state connectivity. We calculated how accurate the predictions are at replicating the observed patterns of the MEG-measured functional connectivity. In order to unravel the richness of the whole-scan connectivity that results from our MEG recordings, we used non-negative matrix factorization to decompose the MEG-measured functional connectivity into fundamental components, and calculated how accurately those components are predicted by the algorithms of Gõni et al. (2014) applied to our structural data. In contrast to previous studies, and because our long-term goal is that similar work will be used in the development of biomarkers, we work with participant-specific connectomes, and only use group-averaged ones to compare our work with existing literature. The methodology is described in Sec. Methods and the results are presented in Sec. Results. The implications of our results are discussed in Sec. Discussion.

## 2 Methods

All analyses were performed using MATLAB (MATLAB and Statistics Toolbox Release 2015a, The MathWorks, Inc., Massachusetts, United States), unless otherwise stated. An outline of the analysis pipeline is shown in Fig. 1.

**Figure 1:**
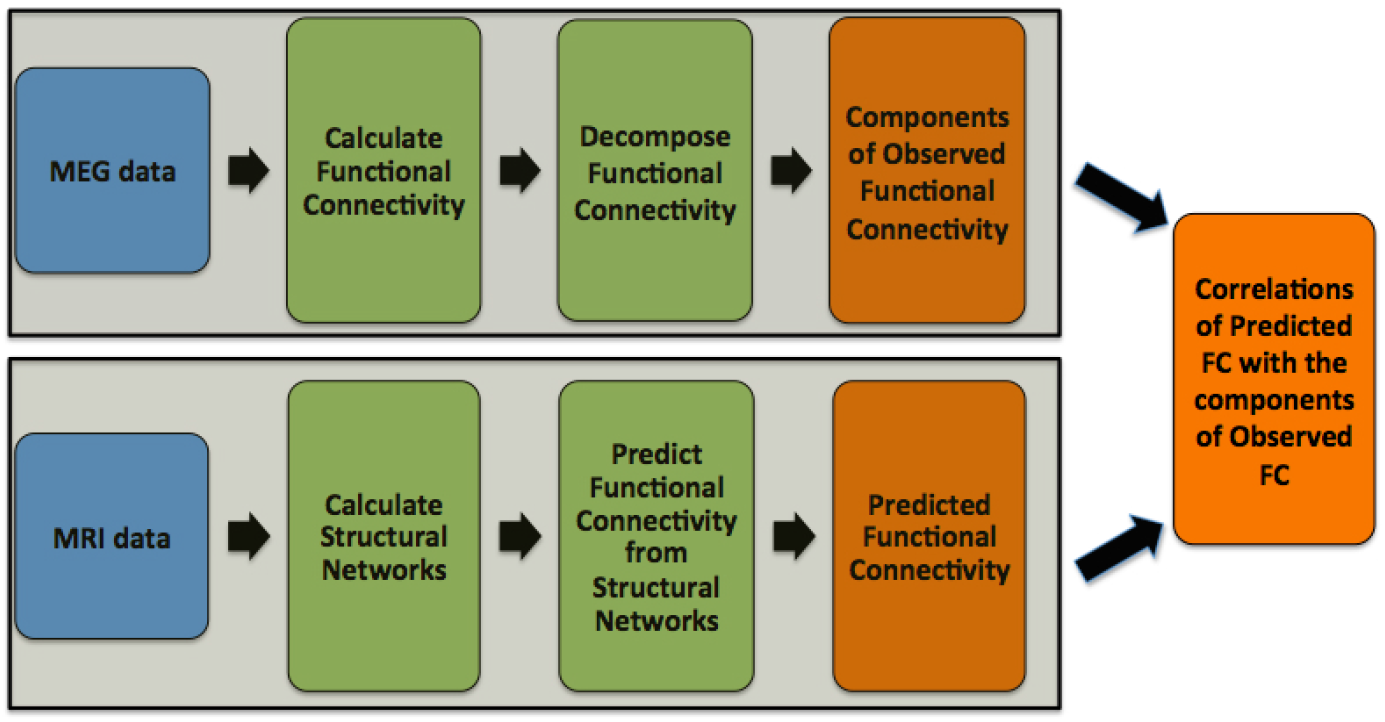
Outline of the methodology.

### 2.1 Data acquisition and preprocessing

Ninety healthy participants (58 females, age: 19 — 34 years, mean age: 23.7 years, SD: 3.4 years) were scanned at the Cardiff University Brain Research Imaging Centre (CUBRIC). This is a subset of the participants from the “100 Brains” and UK MEG Partnership scanning projects (Godfrey and Singh, 2020), and includes the participants who successfully went through the scans described below. All procedures were given ethical approval by the Cardiff University School of Psychology Ethics Committee, and all participants gave written informed consent before taking part.

MRI was carried out on a GE Signa HDx 3T scanner (GE Healthcare, Milwaukee, WI). T1-weighted structural data were acquired using an axial three-dimensional fast spoiled gradient recalled sequence with the following parameters: TR = 8 ms, TE = 3 ms, TI = 450 ms; flip angle = 20°; voxel size = 1 mm; field of view (FOV) ranging from 256 × 192 × 160 mm^3^ to 256 × 256 × 256 mm^3^ (anterior-posterior / left-right / superior-inferior). The T1 images were down-sampled to 1.5 mm isotropic resolution.

Diffusion-weighted MRI data were acquired using a peripherally cardiac-gated sequence with b = 1200 s/mm^2^, TR = 20 s, TE = 90 ms, isotropic resolution of 2.4 mm, zero slice gap, FOV = 230 mm. Data were acquired along 30 unique and isotropically distributed gradient orientations. Three images with no diffusion weighting were also acquired. The diffusion images were co-registered to the T1-weighted images and corrected for head movement and eddy current distortions. Free-water correction was also performed.

The acquisition and preprocessing of mcDESPOT data (Deoni et al., 2008) was done as described by Zacharopoulos et al. (2017). We briefly describe the method here. The acquisition consisted of spoiled gradient recall (SPGR) images for eight flip angles, one inversion recovery SPGR (IR-SPGR) and steady-state free precession (SSFP) images for eight flip angles and two phase-cycling angles. Twenty-five images were acquired for each participant. All images were acquired in sagittal orientation with a slice matrix of 128 × 128(1.72 × 1.72 mm resolution) with a minimum of 88 slices (slice thickness =1.7 mm). Additional slices were acquired for some participants, in order to ensure full head coverage. The parameters used for each sequence were: SPGR: TE = 2.112 ms, TR = 4.7 ms, flip angles = 3°, 4°, 5°, 6°, 7°, 9°, 13° and 18°. IR-SPGR: TE = 2.112 ms, TR = 4.7 ms, IR = 450 ms, flipangle = 5°. SSFP: TE = 1.6 ms, TR = 3.2 ms, flip angles of 10.59°, 14.12°, 18.53°, 23.82°, 29.12°, 35.29°, 45° and 60°, and phase-cycling angles of 0° and 180°. All images were linearly coregistered to the 13° SPGR image to correct for subject motion. Non-brain tissue was removed using a mask computed with the BET algorithm (Smith et al., 2002). Registration and brain masking were performed with FSL (http://www.fmrib.ox.ac.uk/fsl/, Woolrich et al. (2009); Smith et al. (2004); Jenkinson et al. (2012)). The images were then corrected for B1 inhomogeneities and off-resonance artifacts, using maps generated from the IR-SPGR and 2 phase-cycling SSFP acquisitions, respectively. The 3-pool mcDESPOT algorithm was then used to identify the fast (water constrained by myelin) and slow (free-moving water in intra- and extra-cellular space) components of the T1 and T2 times, and a non-exchanging free-water component (Deoni et al., 2013). The fast volume fraction was taken as a map of the myelin water fraction. The myelin volume fraction was calculated as the ratio of myelin-bound water to total water.

Five-minute whole-head MEG recordings were acquired in a 275-channel CTF radial gradiometer system, at a sampling rate of 1200 Hz. Twenty-nine additional reference channels were recorded for noise cancellation purposes and the primary sensors were analysed as synthetic third-order gradiometers (Vrba and Robinson, 2001). Participants were seated upright in a magnetically shielded room with their head supported with a chin rest to minimise movement. They were asked to rest with their eyes open and to fixate on a central red point, presented on either a CRT monitor or LCD projector. Horizontal and vertical electro-oculograms (EOG) were recorded to monitor eye blinks and eye movements. Recordings were also acquired while the participants performed tasks after the completion of the resting-state recording, but those recordings were not used in the analysis presented here. To achieve MRI/MEG co-registration, fiduciary markers were placed at fixed distances from three anatomical landmarks identifiable in the participant’s T1-weighted anatomical MRI scan, and their locations were manually marked in the MR image. Head localization was performed at the start and end of the MEG recording. The data were subsequently pre-processed in a manner identical to that described in Koelewijn et al. (2019). Specifically, all datasets were down-sampled to 600 Hz, and filtered with a 1 Hz high-pass and a 150 Hz low-pass filter. The datasets were then segmented into 2s epochs, and were visually inspected. Epochs exhibiting large head movements, muscle contractions, or ocular artefacts were excluded from subsequent analysis.

### 2.2 Network construction

The Automated Anatomical Labeling (AAL) atlas (Tzourio-Mazoyer et al., 2002) was used to define, for each participant, the 90 cortical and sub-cortical areas of the cerebrum that correspond to the nodes of the structural and functional networks. The correspondence between node numbers and brain areas for the AAL atlas is given in Table 1. Matching between the cortical and sub-cortical areas of each participant and the AAL atlas was performed in ExploreDTI-4.6.8 (Leemans et al., 2009). Each network can be represented as a 90 × 90 symmetric matrix, with each entry indicating the strength of the connection (structural or functional) between the respective nodes. Diagonal elements of those matrices, i.e. self-connections, were set to zero.

**Table 1:** AAL atlas areas.

|  |  |  |  |  |  |  |  |  |  |
| --- | --- | --- | --- | --- | --- | --- | --- | --- | --- |
| 1 | Precentral L | 19 | Supp Motor Area L | 37 | Hippocampus L | 55 | Fusiform L | 73 | Putamen L |
| 2 | Precentral R | 20 | Supp Motor Area R | 38 | Hippocampus R | 56 | Fusiform R | 74 | Putamen R |
| 3 | Frontal Sup L | 21 | Olfactory L | 39 | ParaHippocampal L | 57 | Postcentral L | 75 | Pallidum L |
| 4 | Frontal Sup R | 22 | Olfactory R | 40 | ParaHippocampal R | 58 | Postcentral R | 76 | Pallidum R |
| 5 | Frontal Sup Orb L | 23 | Frontal Sup Medial L | 41 | Amygdala L | 59 | Parietal Sup L | 77 | Thalamus L |
| 6 | Frontal Sup Orb R | 24 | Frontal Sup Medial R | 42 | Amygdala R | 60 | Parietal Sup R | 78 | Thalamus R |
| 7 | Frontal Mid L | 25 | Frontal Med Orb L | 43 | Calcarine L | 61 | Parietal Inf L | 79 | Heschl L |
| 8 | Frontal Mid R | 26 | Frontal Med Orb R | 44 | Calcarine R | 62 | Parietal Inf R | 80 | Heschl R |
| 9 | Frontal Mid Orb L | 27 | Rectus L | 45 | Cuneus L | 63 | SupraMarginal L | 81 | Temporal Sup L |
| 10 | Frontal Mid Orb R | 28 | Rectus R | 46 | Cuneus R | 64 | SupraMarginal R | 82 | Temporal Sup R |
| 11 | Frontal Inf Oper L | 29 | Insula L | 47 | Lingual L | 65 | Angular L | 83 | Temporal Pole Sup L |
| 12 | Frontal Inf Oper R | 30 | Insula R | 48 | Lingual R | 66 | Angular R | 84 | Temporal Pole Sup R |
| 13 | Frontal Inf Tri L | 31 | Cingulum Ant L | 49 | Occipital Sup L | 67 | Precuneus L | 85 | Temporal Mid L |
| 14 | Frontal Inf Tri R | 32 | Cingulum Ant R | 50 | Occipital Sup R | 68 | Precuneus R | 86 | Temporal Mid R |
| 15 | Frontal Inf Orb L | 33 | Cingulum Mid L | 51 | Occipital Mid L | 69 | Paracentral Lobule L | 87 | Temporal Pole Mid L |
| 16 | Frontal Inf Orb R | 34 | Cingulum Mid R | 52 | Occipital Mid R | 70 | Paracentral Lobule R | 88 | Temporal Pole Mid R |
| 17 | Rolandic Oper L | 35 | Cingulum Post L | 53 | Occipital Inf L | 71 | Caudate L | 89 | Temporal Inf L |
| 18 | Rolandic Oper R | 36 | Cingulum Post R | 54 | Occipital Inf R | 72 | Caudate R | 90 | Temporal Inf R |

The white matter (WM) tracts linking the brain areas are the connections, or edges, of the structural networks. The first step in the construction of the structural networks was to perform tractography using a deterministic streamline algorithm in MRtrix 3.0 (Tournier et al., 2019; Dhollander et al., 2016, 2019) (function `tckgen SD_stream`). WM tracts were seeded in the white matter, using a WM mask generated from the T1-weighted images using `FSL fast` (Smith et al., 2004; Jenkinson et al., 2012; Zhang et al., 2001). The response function was calculated with spherical harmonic order l_max_ = 6. The maximum allowed angle between successive steps was 45°, and the minimum and maximum tract lengths were 30 and 250 mm respectively. The algorithm was set to select 2 × 10^5^ streamlines.

The tractograms generated were used to identify the structural connectomes of the participants, using the MRtrix function `tck2connectome`. Tracts that had been generated with fewer than 5 streamlines were excluded from the structural connectome construction. Three structural networks were derived to represent the structural connectome for each participant. The edges of those networks were weighted by the number of streamlines scaled by the volume of the two connected nodes (NS), the mean fractional anisotropy (FA) along the streamlines (which has been shown to be correlated with myelination (de Santis et al., 2014), in single-tract voxels), and the mean radial diffusivity (RD) of the streamlines (which has been shown to correlate with axonal diameter by Barazany et al. (2009)). The `tcksample` function was again used to scan the myelin volume fraction (MVF) images, assign the proportion of the MVF that each streamline is responsible for and divide by the tract length. Then, for each tract, the average myelin measure (MM) along its streamlines was defined to be its edge weight, in order to create a fourth structural network for each participant (MM-weighted). We note that a different myelin measure, the g-ratio, has previously been used as an edge weight in structural connectomes (Mancini et al., 2018). The matrices that resulted from these four edge-weightings were normalized by dividing by the largest value of each matrix, so that, within any given matrix, the values range from 0 to 1. In order to assess the predictive capability of combinations of the WM attributes, we derived data-driven combinations of them based on the method described by Dimitriadis et al. (2017), resulting in composite structural connectivity matrices, and used them as edge-weights in the SC matrices used to predict the functional connectivity (FC). Finally, in order to have a measure of the impact of these four edge weightings on our analyses, we also constructed a binarized network for each participant, in which the strength of a structural connection was set to 1 if a WM tract linking the corresponding brain areas existed, and it was set to 0 if such a white matter tract did not exist. We use ‘b’ to denote those binarized networks.

The modular organisation of the structural networks and any differences in their hubs are also of relevance to the analysis. In order to develop some understanding of the differences between structural networks that have been weighted with the different metrics, we calculated the node degree and the betweenness centrality of each node, normalised them, and averaged them to calculate a hub score for each node, in a manner similar to that described by Betzel et al. (2014). This procedure was repeated for the structural networks of each participant, and for each of the structural edge weightings used in our analysis. The nodes with the 10 highest hub scores were identified as hubs. We also calculated the modular structure of each network for each participant using the Brain Connectivity Toolbox.

The functional networks were constructed in a manner similar to that described by Koelewijn et al. (2019). Specifically, the MEG sensor data were source-localised using FieldTrip (`RRID: SCR_004849`) version 20161011 (Oostenveld et al., 2011), with an LCMV beamformer on a 6 mm grid, using a single-shell forward model (Nolte, 2003), where the covariance matrix was constructed in each of four frequency bands: delta (1 — 4 Hz), theta (3 — 8 Hz), alpha (8 — 13 Hz) and beta (13 — 30 Hz). For each band, the beamformer weights were normalized using a vector norm (Hillebrand et al., 2012), data were normalized to the MNI template, and mapped to 90 nodes based on the Automatic Anatomical Labelling (AAL) atlas (Tzourio-Mazoyer et al., 2002). Epochs were concatenated to generate a continuous virtual-sensor time course for each voxel and then band-passed into the above-mentioned frequency bands. For each of the 90 AAL regions, the virtual channel with the greatest temporal standard deviation was selected as the representative one.

The resulting 90 time series were orthogonalized in order to avoid spurious correlations, using symmetric orthogonalization (Colclough et al., 2015). A Hilbert transform was then used to obtain the oscillatory amplitude envelope. The data were subsequently de-spiked using a median filter in order to remove artifactual temporal transients, down-sampled to 1 Hz, and trimmed to avoid edge effects (removing the first two and the last three samples). Amplitude correlations were calculated by correlating the 90 down-sampled Hilbert envelopes to each other, and were converted to variance-normalized z-scores by applying a Fisher transform. This choice was motivated by the fact that such correlations have been shown to be one of the most robust and repeatable electrophysiological connectivity measures (Colclough et al., 2016; Godfrey and Singh, 2020). We adjusted each participant’s FC matrix to correct for global session effects (Siems et al., 2016). These effects can be generated by experimental confounds such as head size, head motion and position within the MEG helmet. Such correction procedures are common in fMRI analyses, although it is still not clear what is the best method for post-hoc standardization (Yan et al., 2013). We adopted a variant of z-scoring, in which the null mean and standard deviation of connectivity is estimated by fitting a Gaussian (Lowe et al., 1998) to the noise peak (± 1 standard deviation) of the distribution. This estimated mean and standard deviation were then used to z-score each Fisher’s z connectivity value for that participant. Finally, the weakest 80% of values were removed. We note that we only considered positive amplitude-amplitude correlations in our analysis, because the few negative correlations that resulted were very faint and not robust, and therefore amount to noise in our experiment. Robust negative correlations are generally not seen with static amplitude-amplitude correlations in MEG experiments.

Functional connectivity derived from fMRI data has been shown to be impacted by the Euclidean distance between brain areas (for example Alexander-Bloch et al. (2012); Gõni et al. (2014)). In order to assess whether that is the case in our data, the Euclidean distance was plotted against the functional connectivity strength.

### 2.3 Decomposition of observed functional connectivity

The whole-scan MEG resting-state functional connectivity is complex, and could potentially be broken up into fundamental parts. Non-negative matrix factorization (NMF) (Lee and Seun, 1999) is a mathematical method that allows a set of matrices to be decomposed into their fundamental components. It has been previously used in MEG studies to decompose time-dependent FC (Phalen et al., 2019), and allowed for differences between Schizophrenia patients and healthy controls to be quantified. In this work, NMF was used to derive the components that comprise the observed FC (FC_o_) for the group of participants, and to identify the contribution that each component has on the whole-scan connectivity of each participant. In NMF, the number of components for the decomposition needs to be specified a-priori, and the resulting components depend on that pre-specified number. This needs to be done with care: a small number of components can result in some details of the FC_o_ not being captured, while a large number of components can result in at least some of the components being relevant to only a very small number of participants of the group. In our analysis, the number of components was chosen to be the maximum number for which all components were present in at least half of the total number of participants, with a contribution that is at least 5% of the maximum contribution, ensuring that the component is important in explaining the whole-scan FC_o_ of those participants. The NMF analysis was performed for each frequency band independently, and there was no requirement for the number of components to be identical for the four frequency bands.

As a result of the NMF decomposition, the FC_o_ of each participant (index *i*) for the frequencyband *f* can be written as a sum of the N_*f*_ components FC_*j*_, namely:

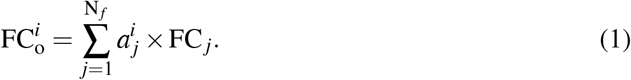

The coefficients *a^i^_j_* are specific to each participant for each frequency band, while the components FC_*j*_ are frequency-band specific.

The motivation for decomposing the FC_o_ in the context of our work, which aims to predict the FC from the SC, comes from the fact that the FC derived from our MEG recordings pertains to the whole 5-minute scan and is very complex. At the same time, the function-predicting algorithms are powerful but rely on simple interactions between brain areas. Our hypothesis is that the function-predicting algorithms would be better at capturing the components than at capturing the total FC_o_.

Various algorithms can be used to predict the FC given a substrate of structural connections between the cortical and sub-cortical areas. A number of such algorithms were described by Gõni et al. (2014) and implemented in the Brain Connectivity Toolbox (Rubinov and Sporns, 2010). These algorithms calculate the potential predictors of FC based on a structural connectivity (SC) matrix, using different methods to identify the optimal links between brain areas, and then use regression to generate a predicted FC that best matches the observed FC. They favor structural connections that are stronger, and take into account both direct and indirect connections between brain areas.

In this work, the search-information (SI) and the shortest-path-length (SPL) algorithms were used to predict the FC for each participant using the five different structural edge-weightings described earlier, with the Euclidean distance also used as a predictor in those algorithms. The analysis was performed for each participant and each frequency band independently. The inverse transform was used to convert the edge weights to distances. The predicted FC was calculated both for the total FC_o_, and for each of the components derived through NMF. In order to assess the impact of the structural edge-weightings and the function-predicting algorithms on the predicted FC (FC_p_), the Pearson correlations between FC_p_ derived with all possible edgeweighting - algorithm pairs were calculated, at participant-level for the total (pre-NMF) FC_o_. In order to assess the reliability of the predictions derived with the SI and SPL algorithms for our electrophysiologically-measured FC, the correlations between the FC_o_ and the FC_p_ were calculated, both for the total connectivity (pre-NMF) and for the NMF-derived components. Because we are interested in the reliability with which the functional connections that were realised are predicted, we do not include, in the correlation calculation only, the predicted functional connections that were zero in the FC_o_ matrices of the participants.

Finally, in order to compare our results with results that exist in the literature, we calculated the participant-average (group-average) structural connectomes, using the edges that show up in at least 10% of the participants, for each of the five edge weightings. We also calculated the group-average functional connectomes for FC_o_, using the edges that show up in at least 10% of the participants, for each of the four frequency bands. We used applied the SPL and SI algorithms on the average structural connectomes to derive the FC_p_, and calculated the correlations between FC_p_ and FC_o_ as before.

## 3 Results

### 3.1 Observed functional connectivity

The strength of the connections of FC_o_ depended on the Euclidean distance between brain areas (Fig. 2), a result that is in agreement with what has been observed for fMRI functional connectivity (for example by Gõni et al. (2014) and by Alexander-Bloch et al. (2012) - also note a discussion of the relationship between MEG-derived FC and Euclidean distance in Tewarie et al. (2019)). There were more and stronger functional connections between brain areas that are directly linked with a WM tract (Fig. 2). However, calculating the mean Euclidean distance between brain areas which are linked with a WM tract and that of those which are not we observe that more functional connections exist between brain areas that are close to each other, and therefore have smaller Euclidean distance (Table 2). The distributions of the number of FC_o_ connections versus the Euclidean distance for linked and unlinked brain areas is shown in Fig. 3. The apparent dependence of the strength of functional connectivity on whether or not the brain areas are linked with a WM tract could be due to the fact that, by construction, brain areas that are closer to each other are more likely to have a direct WM tract assigned between them.

**Figure 2:**
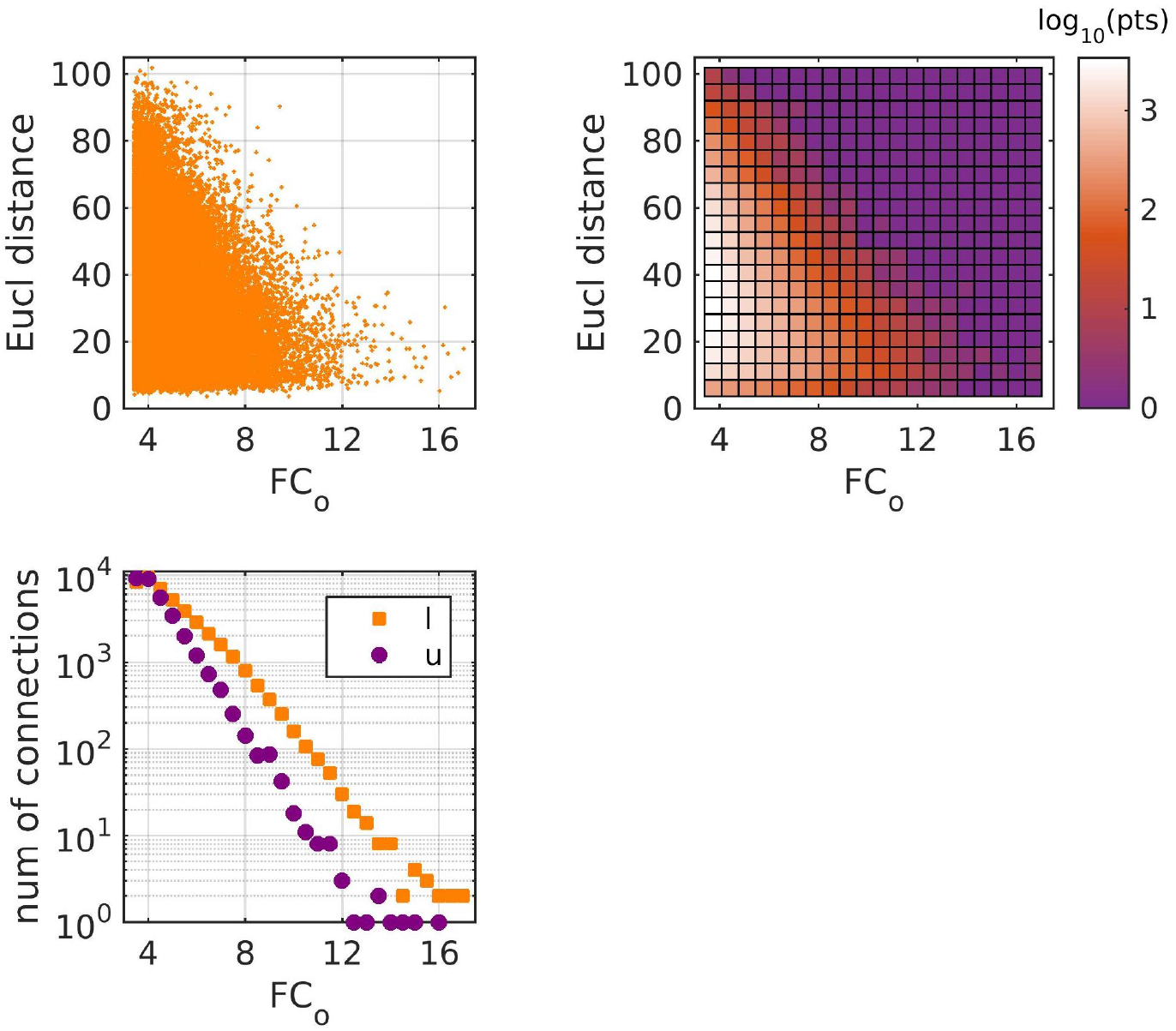
Top left: Euclidean distance vs FC_o_ strength between brain areas. Top right: Density plot of the top left plot, where the color in each square represents the log_1_θ of the number of points in that square. Bottom: Number of connections vs functional connection strength, for brain areas that are linked/unlinked with a WM tract. All three plots refer to the z-scored FC_o_ and contain all connections from all 90 participants.

**Figure 3:**
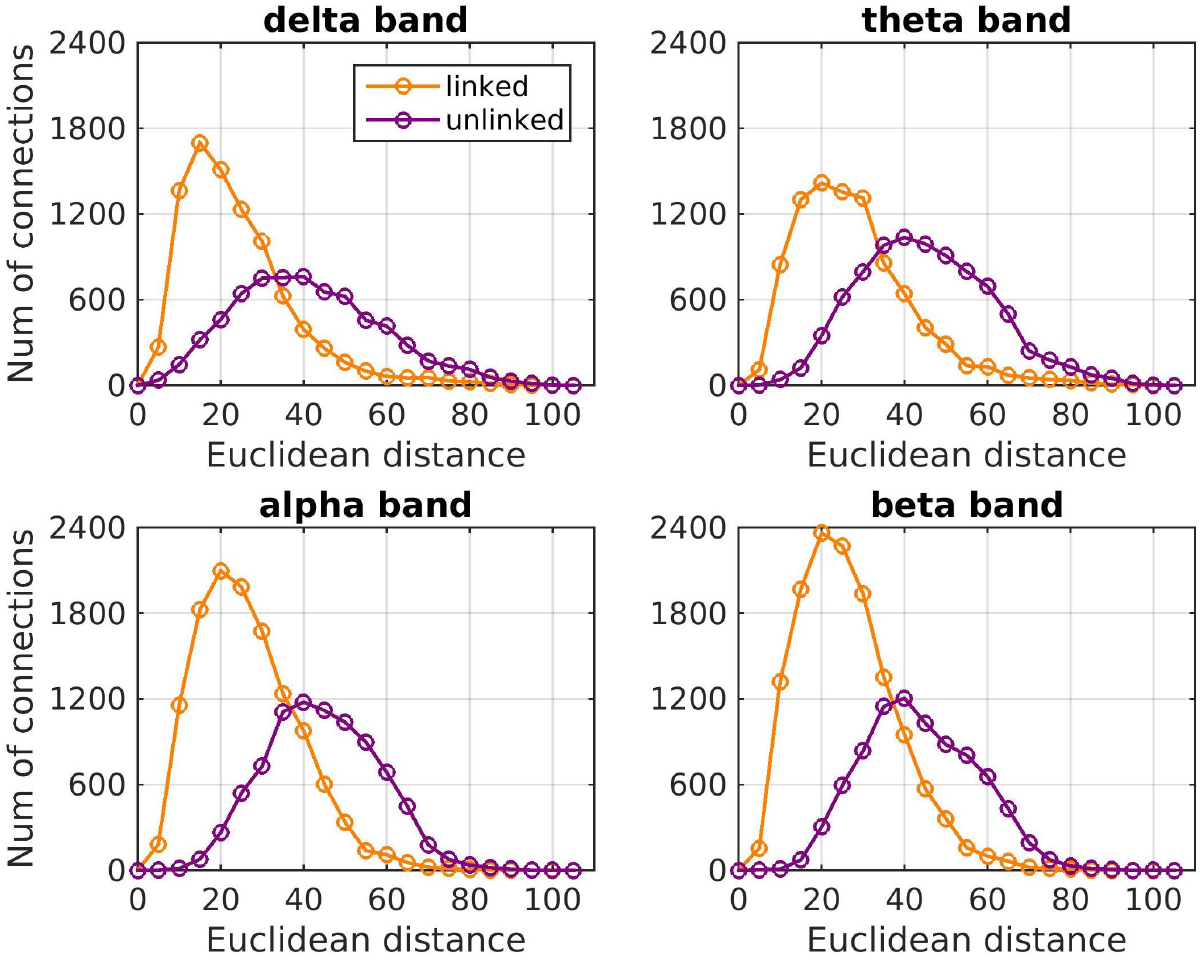
Histogram of the number of FC_o_ connections versus the Euclidean distance (in mm) for brain areas that are linked or unlinked with a WM tract, for each frequency band.

**Table 2:** Mean Euclidean distance (mm) between brain areas that exhibit functional connectivity in each frequency band, for areas that are linked with a WM tract, and those that are not linked.

| Frequency band | Mean Euclidean distance |  |
| --- | --- | --- |
|  | Linked | Unlinked |
| $\delta$ | 23.9 | 41.0 |
| $\theta$ | 27.7 | 45.2 |
| $\alpha$ | 26.6 | 44.4 |
| $\beta$ | 26.2 | 43.6 |

This implies that it is the Euclidean distance, rather than the presence or absence of a link between two brain areas, that plays the important role in the strength of the functional connectivity between those brain areas.

Based on the considerations described in Methods regarding the number of components in the NMF algorithm, the number of components depended on the frequency band of the FC_o_. The maximum number of components for which all components were present for at least half of the participants with a contribution that was at least 5% of the maximum contribution, was 8 for the delta band, 10 for the theta band, 9 for the alpha band and 9 for the beta band. These components are shown in Fig. 4, where the strongest 5% of the connections for each FC component are shown. The contribution of the connectivity of each participant to each component is shown in Fig. 5. We discuss the components in detail in the Discussion.

**Figure 4:**
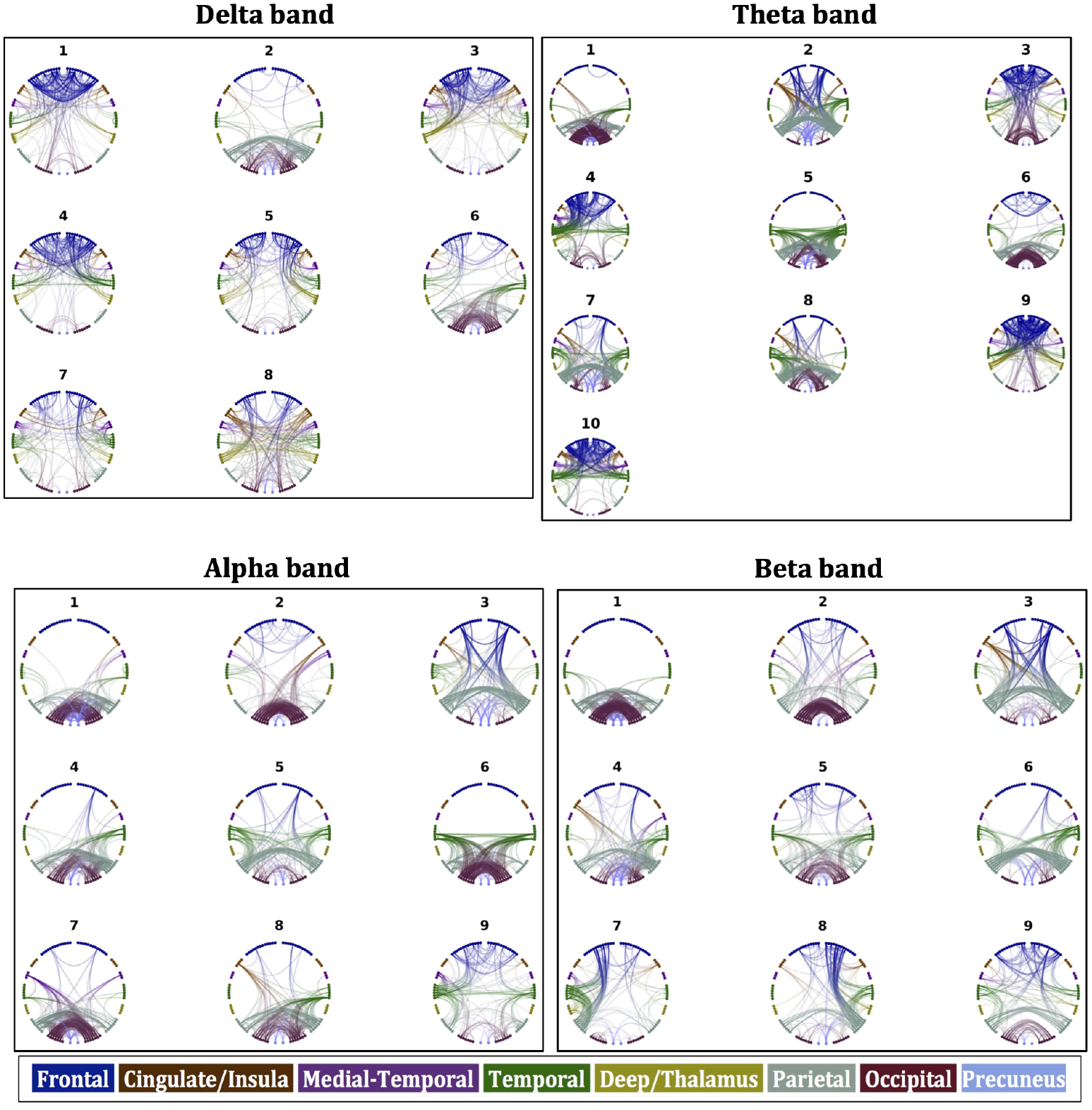
Components of FC_o_ for the four frequency bands.

**Figure 5:**
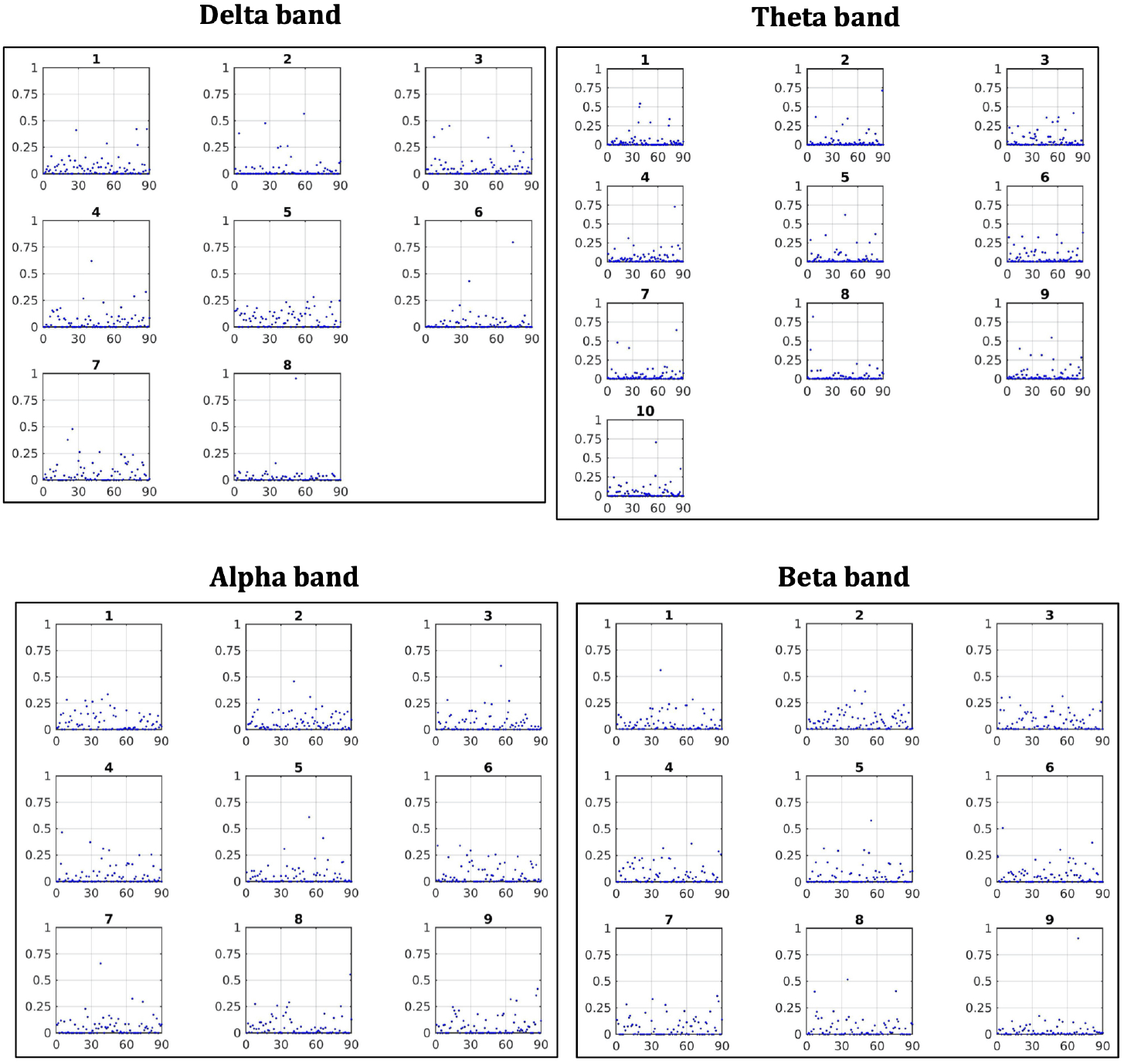
Relative contribution from each participant to each component of the FC_o_, for the four frequency bands. The horizontal axis is over the 90 participants.

### 3.2 Predicted FC

Table 3 shows the correlations between the metrics used as edge weights in the SC matrices, for all tracts and all participants. We stress that the only difference in the 5 SC matrices derived for each participant comes from the edge weights. For each participant, the SC matrices are constructed from the same tracts, but each matrix results from assigning the strength of the structural connectivity of the tracts based on one of the 5 metrics.

**Table 3:** Correlations of metrics across all SC connections and participants.

|  | NS | FA | MM | RD | ED |
| --- | --- | --- | --- | --- | --- |
| NS | 1 | -0.063 | 0.749 | 0.008 | -0.329 |
| FA | -0.063 | 1 | 0.190 | -0.359 | 0.535 |
| MM | 0.749 | 0.190 | 1 | -0.095 | -0.084 |
| RD | 0.008 | -0.359 | -0.095 | 1 | -0.190 |
| ED | -0.329 | 0.535 | -0.084 | -0.190 | 1 |

The lack of strong correlations between the metrics (with the exception of the NS-MM pair) implies a substantial dependence of the results of the analysis on the choice of edge-weight. Fig. 6 shows the SC matrices used in the function-predicting algorithms for one representative participant. Each edge-weighting results in a distinct pattern of SC strength for the structural connections.

**Figure 6:**
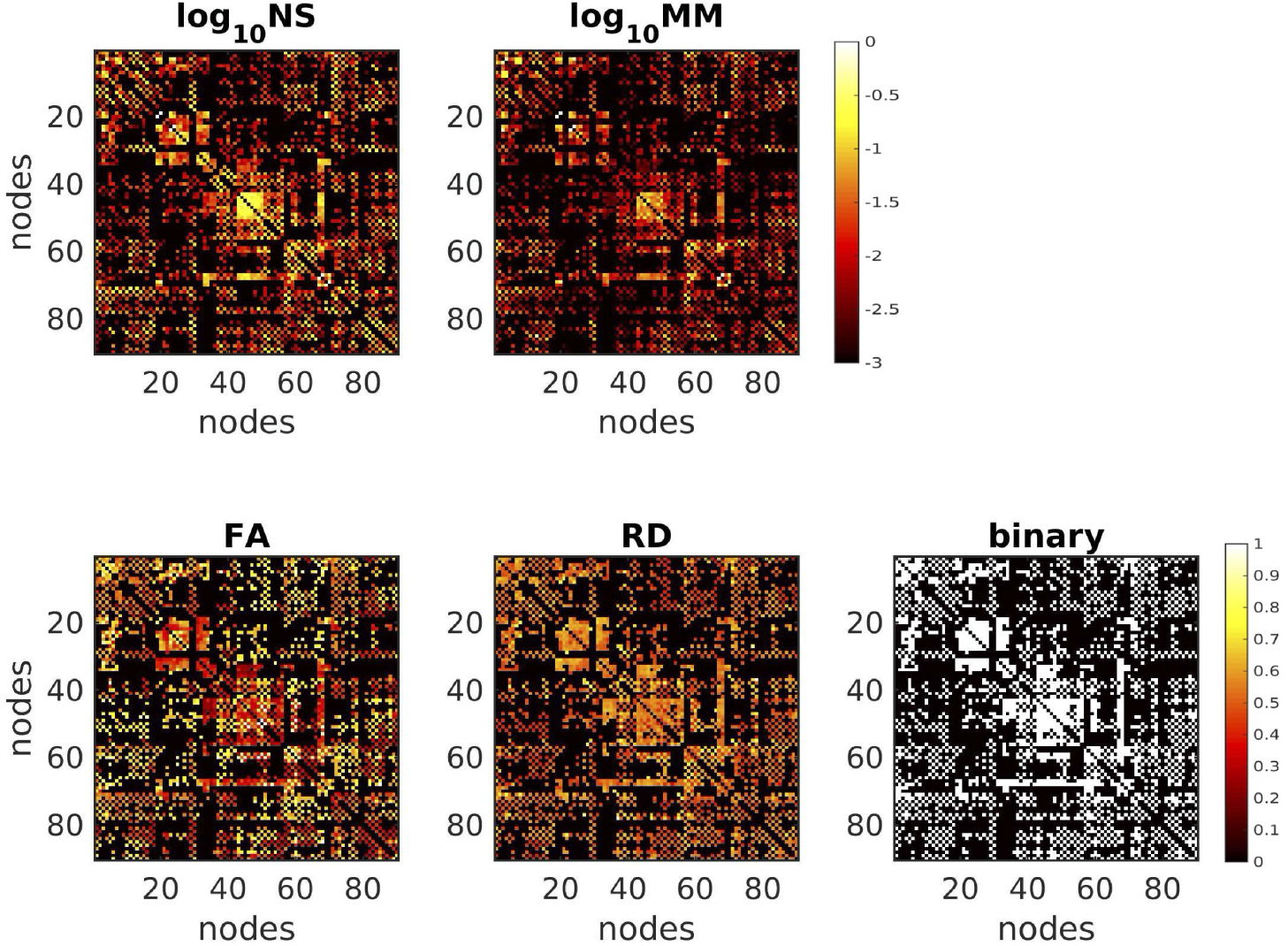
SC matrices for the five different edge-weightings, for one participant. In order to demonstrate the detail of the differences for the NS and MM matrices, we plot the log_10_ of those metrics.

The hub score for the nodes of each SC network depended on the metric used to weight the edges. The left and right precuneus are hubs for the majority of participants regardless of which metric is used as edge weight, with the exception of the binarized graphs. Similarly, the thalamus is a hub for most participants regardless of the metric used as edge weight, with the exception of MM. The caudate is a hub for most participants when the NS or the RD are used. The superior parietal gyrus is a hub when any metric other than NS is used as edge weight. The nodes that are hubs for each participant are shown in Fig. 8. The modularity of the networks depends on the choice of edge weight, as shown in Fig. 7. The MM and NS result in more modular networks than when the other metrics are used. The mean and standard deviation of the number of modules are shown in Table 4.

**Figure 7:**
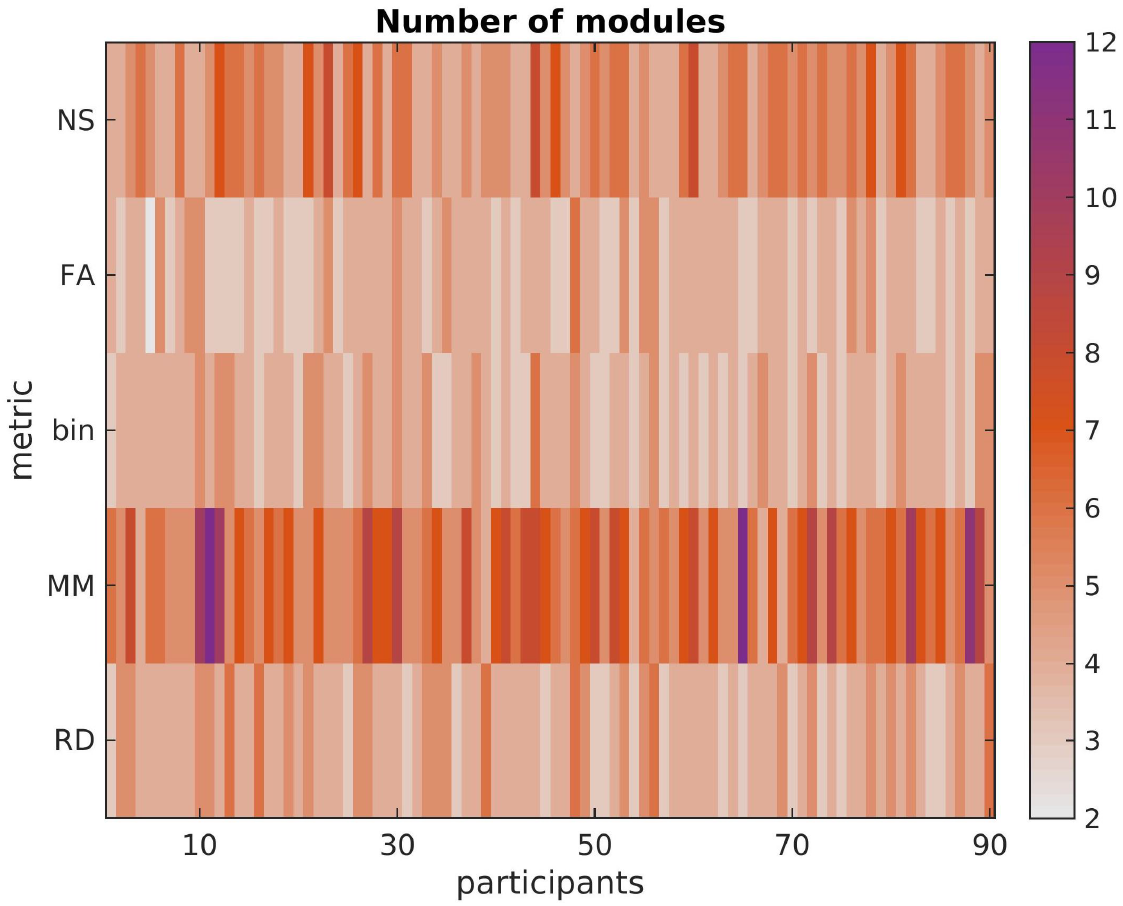
Number of modules in the SC matrices of each participant, for each structural edge-weighting. Using the NS or the MM as edge weights results in a more modular structure than when using the FA, the RD, or the binarized SC matrices.

**Figure 8:**
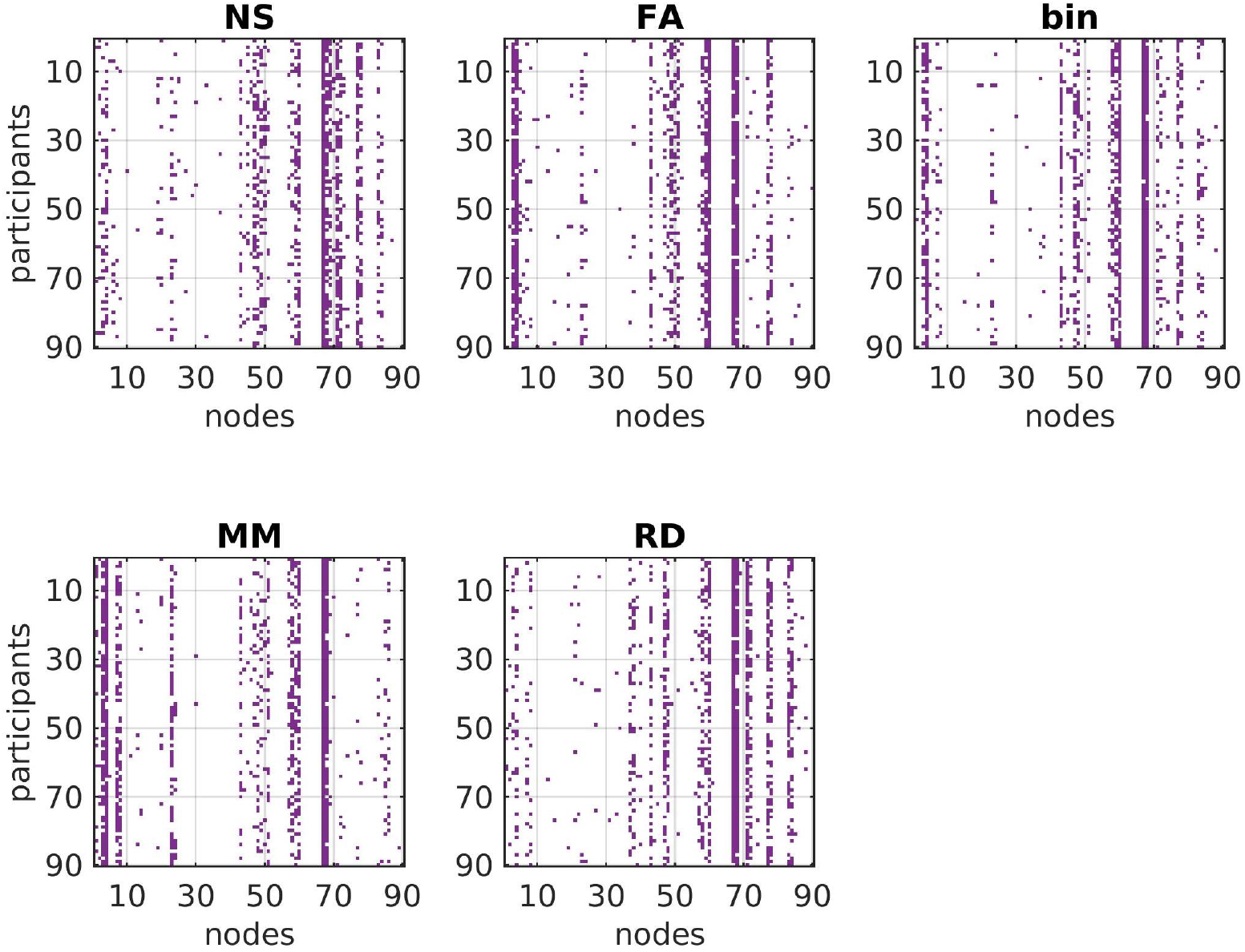
Graphical representation of the fact that the hub nodes for each participant have some similarities and some differences across structural edge-weightings. The brain areas that the number of each node corresponds to are listed in Table 1.

**Table 4:** Mean and standard deviation of the number of modules for the SC matrices.

|  | mean | SD |
| --- | --- | --- |
| NS | 5.16 | 1.07 |
| FA | 3.78 | 0.72 |
| binary | 3.94 | 0.69 |
| MM | 6.47 | 1.74 |
| RD | 4.18 | 0.80 |

The patterns of FC_p_ derived using the different structural edge-weightings and the two FC-predicting algorithms, exhibited high correlations, all with a mean value of above 0.75 (Fig. 9). However, for some participants and some edge-weightings and algorithms the correlations between the FC_p_ were lower, with values that sometimes go below 0.6. It is noteworthy that the correlations were highest for the delta band, regardless of the pair of algorithm/metric used. We note that the composite structural connectomes that were derived with the algorithm of Dimitri-adis et al. (2017), even though very good at capturing differences between populations (Dimitriadis et al. (2017); Clarke et al. (2020)), did not provide any improvement in the relationship between structure and function compared to the single-metric structural connectomes, possibly because the true relationship is more complex than that implied by the linear data-driven algorithm. For that reason we do not discuss them any further.

**Figure 9:**
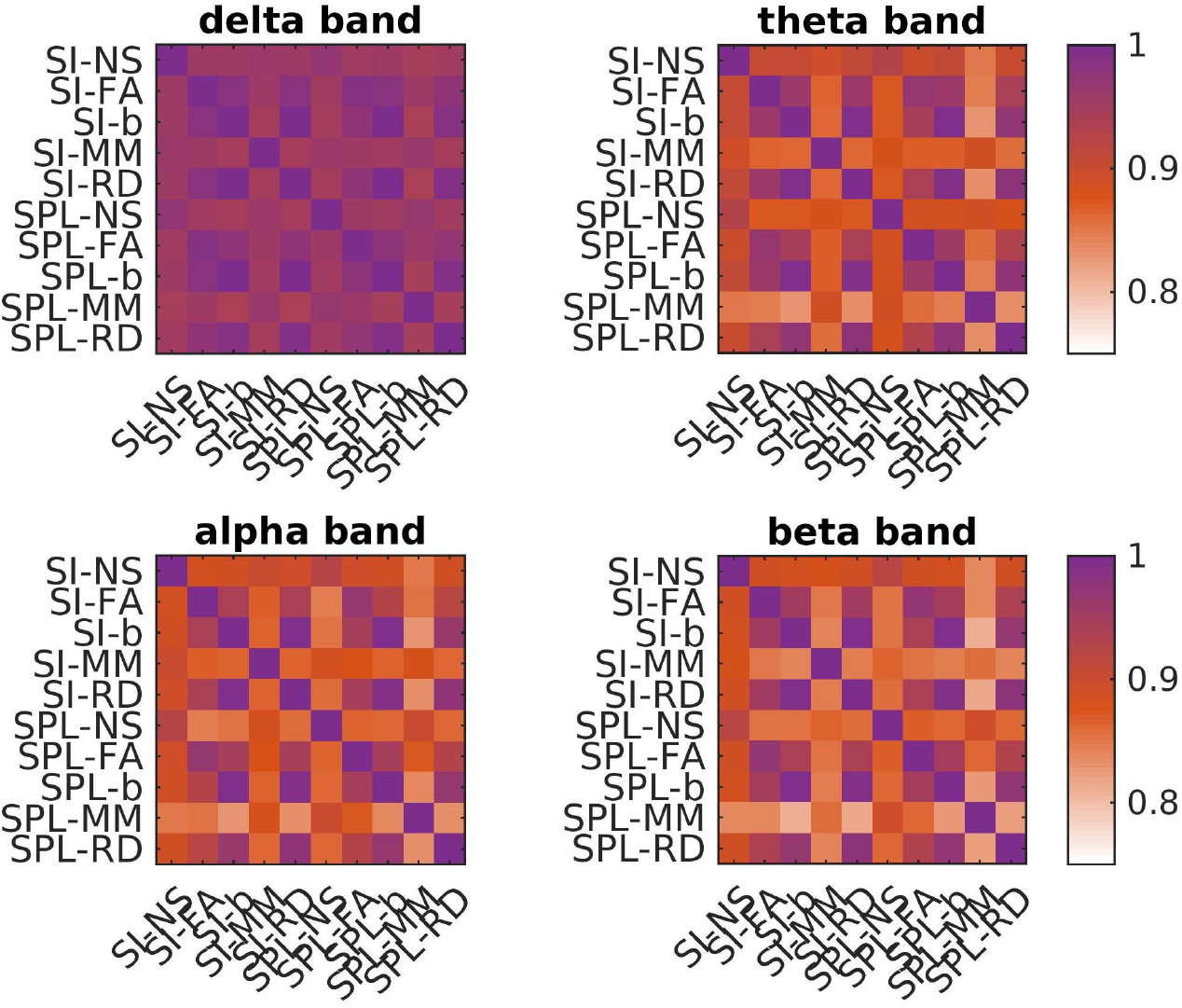
Mean of the correlations of FC_p_ over participants, with different combinations of structural edge-weightings and function-predicting algorithms. SI = search-information, SPL = shortest-path-length.

The mean values of the coefficients for each of the predictors that are calculated from the ED and each of the other metrics, are listed in Tables 5 and 6, for the SPL and SI algorithms respectively. There is no strong dependence of these values on the frequency band or the algorithm considered.

**Table 5:** Mean (over the 90 participants) value of the coefficients that result from the SPL algorithm, when each of the structural metric is used as a predictor with the Euclidean distance, for each frequency band.

| freq. band | NS | ED | FA | ED | bin | ED | MM | ED | RD | ED |
| --- | --- | --- | --- | --- | --- | --- | --- | --- | --- | --- |
| delta | -0.005 | -0.840 | -0.035 | -0.894 | -0.083 | -0.821 | -0.001 | -0.920 | -0.047 | -0.800 |
| theta | -0.006 | -0.551 | -0.046 | -0.632 | -0.097 | -0.549 | -0.002 | -0.672 | -0.052 | -0.532 |
| alpha | -0.015 | -0.838 | -0.069 | -1.034 | -0.197 | -0.852 | -0.005 | -1.101 | -0.011 | -0.817 |
| beta | -0.017 | -0.963 | -0.092 | -1.175 | -0.231 | -0.969 | -0.006 | -1.259 | -0.118 | -0.943 |

**Table 6:**
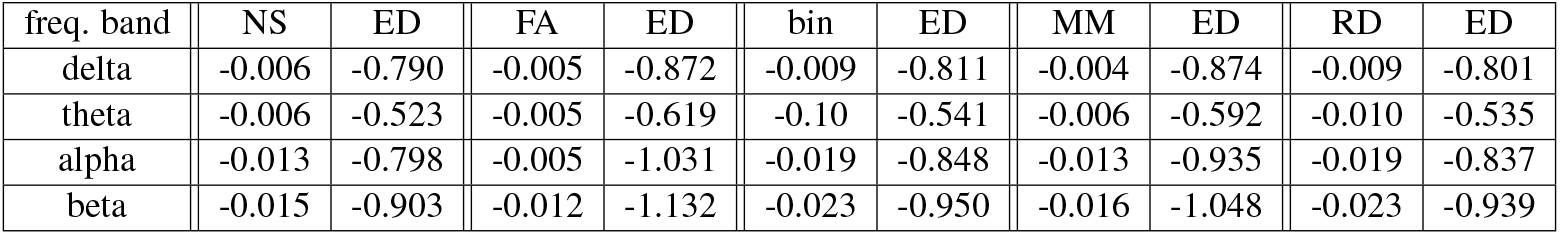
Mean (over the 90 participants) value of the coefficients that result from the SI algorithm, when each of the structural metric is used as a predictor with the Euclidean distance, for each frequency band.

### 3.3 Correlations between predicted and observed FC

For each edge-weighting, the mean of the distributions (over participants) of the correlations between FC_o_ and FC_p_ derived with the SPL algorithm was, in the majority of cases, higher than when derived with the SI algorithm. In Tables 7-10 we give the exact values of the correlations for both the SI and the SPL algorithm for all structural edge-weightings and frequency bands, and we highlight the instances in which it would be beneficial to use the SPL algorithm (bold font, 94 instances) and the few instances in which the SI algorithm gave higher correlations (underlined, 15 instances). For the rest, the mean of the correlations was the same for the SI and SPL algorithms. Given that the SPL algorithm is superior to the SI algorithm, the rest of the paper focuses on FC_p_ derived using the SPL algorithm. The mean of the correlation distributions for the total FC_o_ and its components for the 5 SC edge-weightings and the SPL algorithm are shown in Fig. 10. For all five edge-weightings, the correlations for the individual components were higher than the correlations for the total functional connectivity, the only exceptions being three components in the theta band, one in the alpha band and one in the beta band.

**Figure 10:**
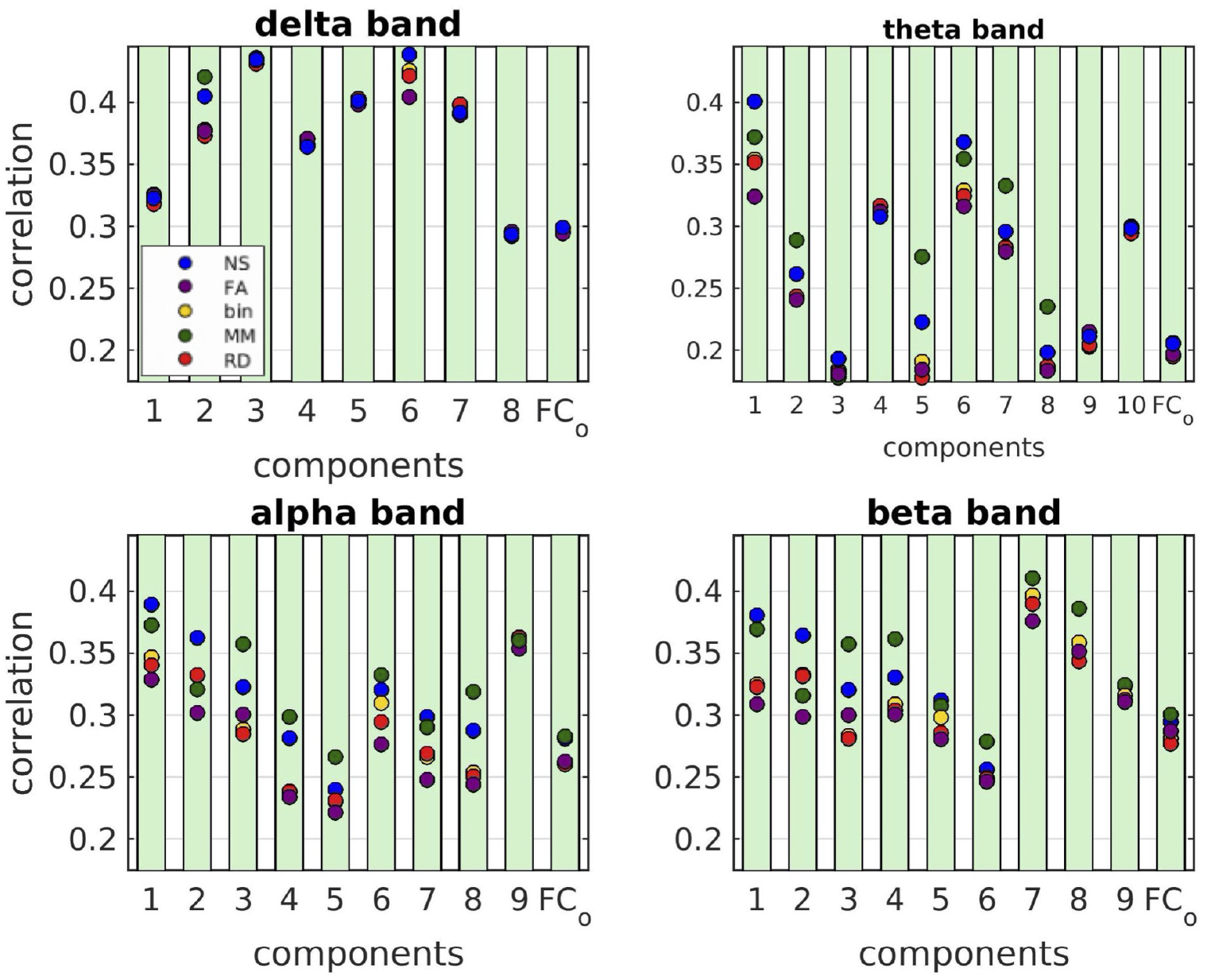
Mean values of the correlations between FC_p_ and FC_o_, for each NMF component and for the total FC_o_ for the 5 structural edge-weightings for the SPL algorithm.

**Table 7:** Mean of the correlations between predicted and observed FC for each metric and algorithm, delta band.

| Components | NS |  | FA |  | binary |  | MM |  | RD |  |
| --- | --- | --- | --- | --- | --- | --- | --- | --- | --- | --- |
|  | SPL | SI | SPL | SI | SPL | SI | SPL | SI | SPL | SI |
| 1 | 0.32 | 0.32 | **0.33** | 0.32 | 0.32 | 0.32 | **0.33** | 0.32 | 0.32 | 0.32 |
| 2 | <u>0.40</u> | 0.41 | 0.40 | 0.40 | 0.40 | 0.40 | 0.40 | 0.40 | 0.40 | 0.40 |
| 3 | 0.43 | 0.43 | 0.43 | 0.43 | 0.43 | 0.43 | 0.44 | 0.44 | 0.43 | 0.43 |
| 4 | <u>0.39</u> | 0.40 | 0.39 | 0.39 | 0.40 | 0.40 | 0.39 | 0.39 | 0.40 | 0.40 |
| 5 | 0.36 | 0.36 | 0.37 | 0.37 | 0.37 | 0.37 | **0.37** | 0.36 | 0.37 | 0.37 |
| 6 | **0.44** | 0.42 | **0.40** | 0.39 | **0.43** | 0.42 | **0.42** | 0.41 | **0.42** | 0.41 |
| 7 | **0.41** | 0.38 | **0.38** | 0.35 | **0.38** | 0.37 | **0.42** | 0.39 | 0.37 | 0.37 |
| 8 | 0.29 | 0.29 | 0.30 | 0.30 | 0.29 | 0.29 | 0.29 | 0.29 | 0.29 | 0.29 |
| FCo | 0.30 | 0.30 | 0.29 | 0.29 | 0.30 | 0.30 | 0.30 | 0.30 | 0.30 | 0.30 |

**Table 8:** Mean of the correlations between predicted and observed FC for each metric and algorithm, theta band.

| Components | NS |  | FA |  | binary |  | MM |  | RD |  |
| --- | --- | --- | --- | --- | --- | --- | --- | --- | --- | --- |
|  | SPL | SI | SPL | SI | SPL | SI | SPL | SI | SPL | SI |
| 1 | **0.26** | 0.25 | 0.24 | 0.24 | 0.24 | 0.24 | **0.29** | 0.28 | 0.24 | 0.24 |
| 2 | 0.31 | 0.31 | 0.31 | 0.31 | **0.32** | 0.31 | 0.31 | 0.31 | 0.32 | 0.32 |
| 3 | **0.21** | 0.20 | 0.21 | 0.21 | 0.20 | 0.20 | 0.21 | 0.21 | 0.20 | 0.20 |
| 4 | 0.19 | 0.19 | **0.18** | 0.17 | 0.18 | 0.18 | 0.18 | 0.18 | <u>0.18</u> | 0.19 |
| 5 | **0.30** | 0.28 | 0.28 | 0.28 | 0.28 | 0.28 | **0.33** | 0.31 | **0.28** | 0.28 |
| 6 | **0.40** | 0.36 | 0.32 | 0.32 | **0.35** | 0.34 | **0.37** | 0.35 | **0.35** | 0.34 |
| 7 | **0.37** | 0.33 | **0.32** | 0.30 | **0.33** | 0.32 | **0.35** | 0.34 | 0.32 | 0.32 |
| 8 | **0.22** | 0.21 | **0.18** | 0.17 | 0.19 | 0.19 | **0.28** | 0.24 | 0.18 | <u>0.19</u> |
| 9 | **0.30** | 0.29 | **0.30** | 0.29 | 0.29 | 0.29 | **0.30** | 0.29 | 0.29 | 0.29 |
| 10 | 0.20 | 0.20 | 0.18 | 0.18 | <u>0.18</u> | 0.19 | **0.24** | 0.21 | 0.19 | 0.19 |
| FCo | **0.21** | 0.20 | **0.20** | 0.19 | **0.20** | 0.19 | 0.21 | 0.21 | 0.19 | 0.19 |

**Table 9:** Mean of the correlations between predicted and observed FC for each metric and algorithm, alpha band.

| Components | NS |  | FA |  | binary |  | MM |  | RD |  |
| --- | --- | --- | --- | --- | --- | --- | --- | --- | --- | --- |
|  | SPL | SI | SPL | SI | SPL | SI | SPL | SI | SPL | SI |
| 1 | **0.36** | 0.33 | 0.30 | 0.30 | **0.33** | 0.32 | 0.32 | 0.32 | **0.33** | 0.32 |
| 2 | **0.39** | 0.35 | **0.33** | 0.31 | **0.35** | 0.33 | **0.37** | 0.35 | **0.34** | 0.33 |
| 3 | **0.32** | 0.29 | **0.28** | 0.25 | **0.31** | 0.29 | **0.33** | 0.29 | 0.29 | 0.29 |
| 4 | **0.30** | 0.27 | 0.25 | 0.25 | **0.27** | 0.26 | **0.29** | 0.26 | **0.27** | 0.26 |
| 5 | **0.29** | 0.26 | **0.24** | 0.23 | 0.25 | 0.25 | **0.32** | 0.26 | **0.25** | 0.24 |
| 6 | <u>0.36</u> | 0.37 | 0.35 | 0.35 | <u>0.36</u> | 0.37 | 0.36 | 0.36 | <u>0.36</u> | 0.37 |
| 7 | **0.32** | 0.29 | **0.30** | 0.29 | 0.29 | 0.29 | **0.36** | 0.32 | 0.28 | 0.28 |
| 8 | **0.28** | 0.25 | 0.23 | 0.23 | 0.24 | 0.24 | **0.30** | 0.26 | **0.24** | 0.23 |
| 9 | 0.24 | 0.24 | 0.22 | 0.22 | 0.23 | 0.23 | **0.27** | 0.26 | 0.23 | 0.23 |
| FCo | **0.28** | 0.27 | **0.26** | 0.25 | **0.26** | 0.26 | **0.28** | 0.27 | 0.26 | 0.26 |

**Table 10:** Mean of the correlations between predicted and observed for each metric and algorithm, beta band.

| Components | NS |  | FA |  | binary |  | MM |  | RD |  |
| --- | --- | --- | --- | --- | --- | --- | --- | --- | --- | --- |
|  | SPL | SI | SPL | SI | SPL | SI | SPL | SI | SPL | SI |
| 1 | **0.36** | 0.33 | **0.30** | 0.29 | 0.33 | 0.33 | 0.32 | 0.32 | 0.33 | 0.33 |
| 2 | **0.38** | 0.33 | **0.31** | 0.30 | **0.33** | 0.31 | **0.37** | 0.32 | **0.32** | 0.31 |
| 3 | **0.32** | 0.28 | **0.30** | 0.29 | 0.28 | 0.28 | **0.36** | 0.31 | 0.28 | 0.28 |
| 4 | 0.26 | 0.26 | 0.25 | 0.25 | 0.25 | 0.25 | 0.28 | 0.28 | 0.25 | 0.25 |
| 5 | **0.33** | 0.32 | **0.30** | 0.29 | **0.31** | 0.30 | **0.36** | 0.33 | 0.30 | 0.30 |
| 6 | **0.32** | 0.31 | 0.31 | 0.31 | 0.32 | 0.32 | **0.32** | 0.31 | <u>0.31</u> | 0.32 |
| 7 | **0.36** | 0.35 | <u>0.35</u> | 0.36 | **0.36** | 0.36 | **0.39** | 0.37 | <u>0.34</u> | 0.36 |
| 8 | <u>0.40</u> | 0.41 | <u>0.38</u> | 0.39 | 0.40 | 0.40 | <u>0.41</u> | 0.42 | <u>0.39</u> | 0.40 |
| 9 | **0.31** | 0.30 | **0.28** | 0.27 | **0.30** | 0.29 | **0.31** | 0.30 | 0.29 | 0.29 |
| FCo | 0.29 | 0.29 | **0.29** | 0.28 | 0.28 | 0.28 | **0.30** | 0.29 | 0.28 | 0.28 |

The NS-weighting and the MM-weighting gave, in most cases, the highest correlations, and the majority of the distributions resulting from different edge-weightings were statistically significantly different from each other. Interestingly, the binarized graphs performed in several cases better than the FA-weighted graphs. The *p*-values for the comparisons of the distributions resulting from all possible combinations of edge-weightings for each component are shown in Tables 11-14. These tables also show the *p*-values that survive multiple comparison correction with the false-discovery-rate (FDR) algorithm (Benjamini and Yekutieli, 2001; Groppe, 2019) in bold.

**Table 11:** p-value resulting from the comparison of the correlation distributions, for each pair of structural edge weightings for the SPL algorithm, in the delta band. Bold numbers indicate the cases for which the distributions were not statistically significantly different.

| Components | SC edge weightings |  |  |  |  |  |  |  |  |  |
| --- | --- | --- | --- | --- | --- | --- | --- | --- | --- | --- |
|  | NS-FA | NS-bin | NS-MM | NS-RD | FA-bin | FA-MM | FA-RD | bin-MM | bin-RD | MM-RD |
| 1 | 4e-04 | 8e-08 | 1e-04 | 2e-10 | 2e-18 | **0.253** | 4e-16 | 7e-15 | 4e-4 | 2e-15 |
| 2 | 2e-08 | 4e-08 | 3e-06 | 1e-07 | **0.348** | 2e-12 | **0.247** | 1e-12 | 8e-03 | 8e-12 |
| 3 | **0.344** | 1e-09 | 0.014 | 1e-08 | 2e-14 | 7e-03 | 5e-12 | 5e-10 | **0.851** | 2e-09 |
| 4 | 9e-11 | 2e-03 | **0.082** | 0.018 | 3e-09 | 4e-06 | 3e-08 | **0.368** | **0.280** | **0.695** |
| 5 | 5e-07 | 0.013 | 2e-08 | 1e-03 | 5e-15 | **0.281** | 2e-14 | 3e-12 | **0.058** | 8e-13 |
| 6 | 4e-11 | 8e-05 | 5e-07 | 7e-06 | 3e-14 | 1e-10 | 6e-09 | **0.196** | 9e-03 | **0.450** |
| 7 | 6e-07 | 6e-13 | 1e-04 | 5e-14 | 3e-17 | 4e-05 | 2e-17 | 2e-15 | 2e-04 | 7e-16 |
| 8 | 7e-04 | 0.017 | 0.015 | 1e-03 | 3e-07 | 0.018 | 3e-07 | 3e-05 | 0.021 | 8e-06 |
| FC<sub>0</sub> | **0.059** | **0.182** | **0.866** | **0.100** | **0.339** | **0.063** | **0.748** | **0.241** | **0.266** | **0.152** |

**Table 12:** p-value resulting from the comparison of the correlation distributions, for each pair of structural edge weightings for the SPL algorithm, in the theta band. Bold numbers indicate the cases for which the distributions were not statistically significantly different.

| Components | SC edge weightings |  |  |  |  |  |  |  |  |  |
| --- | --- | --- | --- | --- | --- | --- | --- | --- | --- | --- |
|  | NS-FA | NS-bin | NS-MM | NS-RD | FA-bin | FA-MM | FA-RD | bin-MM | bin-RD | MM-RD |
| 1 | 8e-14 | 1e-10 | 6e-08 | 2e-10 | 5e-15 | 5e-14 | 8e-10 | 7e-07 | **0.286** | 2e-06 |
| 2 | 9e-09 | 3e-08 | 2e-12 | 4e-07 | 0.013 | 3e-15 | **0.162** | 4e-15 | **0.704** | 2e-14 |
| 3 | 2e-07 | 1e-04 | 1e-09 | 3e-05 | 8e-04 | 5e-03 | **0.155** | 2e-06 | 0.008 | 2e-03 |
| 4 | 1e-04 | 2e-09 | **0.838** | 6e-10 | 1e-06 | 3e-05 | 3e-06 | 5e-09 | 0.054 | 3e-09 |
| 5 | 8e-07 | 1e-05 | 5e-10 | 1e-07 | 8e-04 | 6e-14 | **0.093** | 2e-13 | 2e-05 | 1e-13 |
| 6 | 5e-14 | 9e-13 | 1e-05 | 1e-12 | 2e-11 | 2e-15 | 3e-04 | 1e-13 | 2e-03 | 1e-12 |
| 7 | 4e-07 | 2e-05 | 4e-15 | 1e-04 | 0.010 | 4e-17 | **0.201** | 2e-16 | **0.860** | 6e-15 |
| 8 | 5e-04 | 9e-04 | 1e-09 | 0.009 | **0.184** | 8e-12 | **0.092** | 4e-12 | **0.278** | 6e-11 |
| 9 | **0.103** | 5e-06 | **0.188** | 2e-04 | 3e-11 | **0.878** | 2e-09 | 1e-04 | 0.007 | 0.001 |
| 10 | **0.283** | 0.001 | **0.046** | 0.003 | 3e-07 | **0.372** | 3e-06 | 2e-04 | **0.679** | 4e-04 |
| FC<sub>0</sub> | **0.066** | **0.053** | **0.763** | **0.022** | **0.973** | **0.037** | **0.664** | **0.085** | **0.457** | **0.044** |

**Table 13:** p-value resulting from the comparison of the correlation distributions, for each pair of structural edge weightings for the SPL algorithm, in the alpha band. Bold numbers indicate the cases for which the distributions were not statistically significantly different.

| Components | SC edge weightings |  |  |  |  |  |  |  |  |  |
| --- | --- | --- | --- | --- | --- | --- | --- | --- | --- | --- |
|  | NS-FA | NS-bin | NS-MM | NS-RD | FA-bin | FA-MM | FA-RD | bin-MM | bin-RD | MM-RD |
| 1 | 2e-19 | 6e-16 | 1e-08 | 6e-16 | 2e-16 | 7e-19 | 2e-05 | 4e-13 | 2e-04 | 2e-13 |
| 2 | 1e-23 | 2e-15 | 3e-22 | 1e-13 | 8e-26 | 1e-16 | 4e-20 | 11e-09 | **0.654** | 1e-07 |
| 3 | 2e-09 | 4e-12 | 3e-12 | 3e-11 | 4e-07 | 2e-15 | 3e-05 | 2e-16 | 0.037 | 6e-16 |
| 4 | 3e-10 | 4e-09 | 2e-04 | 9e-09 | 2e-04 | 1e-15 | **0.073** | 3e-15 | **0.719** | 2e-14 |
| 5 | 4e-07 | 0.002 | 2e-11 | 0.010 | 4e-05 | 1e-12 | 8e-04 | 2e-11 | **0.571** | 3e-10 |
| 6 | 9e-09 | 0.049 | 0.021 | 5e-05 | 6e-15 | 5e-14 | 4e-05 | 1e-06 | 2e-07 | 2e-10 |
| 7 | 8e-12 | 9e-09 | 0.041 | 9e-07 | 7e-13 | 3e-15 | 5e-09 | 1e-10 | **0.150** | 2e-07 |
| 8 | 5e-10 | 2e-08 | 3e-08 | 2e-08 | 9e-07 | 3e-16 | 0.044 | 6e-16 | **0.117** | 1e-15 |
| 9 | 1e-09 | 0.022 | **0.338** | **0.193** | 6e-17 | 3e-11 | 9e-14 | 5e-04 | 0.024 | 0.036 |
| FC<sub>0</sub> | 3e-05 | 2e-06 | **0.689** | 9e-08 | **0.847** | 6e-10 | **0.410** | 1e-07 | **0.214** | 1e-08 |

**Table 14:** p-value resulting from the comparison of the correlation distributions, for each pair of structural edge weightings for the SPL algorithm, in the beta band. Bold numbers indicate the cases for which the distributions were not statistically significantly different.

| Components | SC edge weightings |  |  |  |  |  |  |  |  |  |
| --- | --- | --- | --- | --- | --- | --- | --- | --- | --- | --- |
|  | NS-FA | NS-bin | NS-MM | NS-RD | FA-bin | FA-MM | FA-RD | bin-MM | bin-RD | MM-RD |
| 1 | 2e-16 | 6e-15 | 4e-03 | 1e-14 | 5e-12 | 2e-19 | 4e-05 | 6e-18 | **0.277** | 4e-16 |
| 2 | 2e-23 | 7e-15 | 2e-23 | 5e-14 | 1e-25 | 2e-12 | 3e-20 | 8e-13 | **0.217** | 2e-09 |
| 3 | 2e-10 | 6e-16 | 5e-15 | 2e-15 | 2e-14 | 7e-20 | 2e-11 | 1e-21 | 0.049 | 2e-21 |
| 4 | 3e-09 | 1e-06 | 1e-12 | 5e-08 | 3e-07 | 2e-17 | **0.160** | 4e-16 | 1e-03 | 7e-16 |
| 5 | 8e-10 | 6e-05 | **0.077** | 2e-08 | 3e-11 | 7e-12 | 0.027 | 2e-05 | 2e-09 | 3e-09 |
| 6 | 6e-04 | 7e-03 | 4e-11 | 4e-03 | 0.021 | 5e-12 | **0.235** | 3e-12 | **0.543** | 8e-12 |
| 7 | 1e-08 | **0.681** | 2e-07 | 4e-03 | 3e-14 | 2e-12 | 7e-10 | 2e-06 | 5e-09 | 1e-09 |
| 8 | 0.022 | **0.900** | 1e-11 | 1e-05 | 1e-04 | 6e-11 | 0.007 | 4e-10 | 1e-10 | 4e-13 |
| 9 | 1e-06 | 6e-04 | **0.558** | 7e-06 | 1e-04 | 7e-09 | **0.303** | 3e-05 | 2e-05 | 9e-07 |
| FC<sub>0</sub> | **0.079** | 4e-04 | **0.098** | 1e-05 | 0.016 | 9e-05 | 0.002 | 7e-07 | 0.016 | 1e-08 |

In Fig. 11 we show the distributions (over participants) of the correlation coefficients for each metric and each component, in each of the four frequency bands.

**Figure 11:**
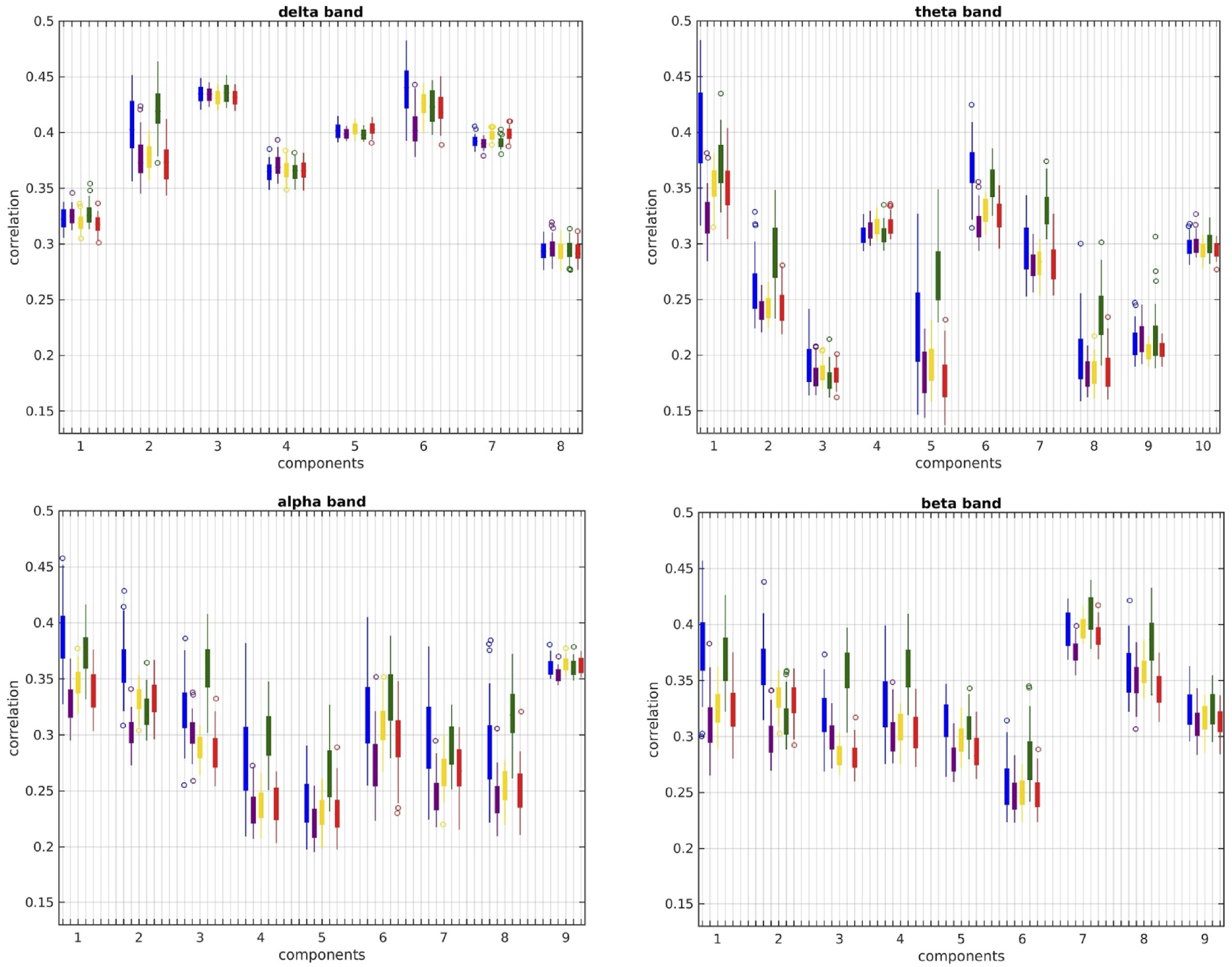
Distributions (over participants) of the correlation coefficients between the observed and the predicted FC NMF components, for each structural edge weighting and frequency band, for the SPL algorithm. The bottom and top ends of the thicker part of each bar indicate the 25^th^ and 75^th^ percentiles respectively, while the whiskers extend to the points that are not considered outliers of each distribution. Any outliers are plotted as circles.

### 3.4 Correlations using group-average connectomes

The correlations that resulted by using group-averaged connectomes are shown in Fig. 12 for the two function-predicting algorithms, for each SC edge-weighting and frequency band. The NS and MM edge-weightings resulted in higher correlations than the other three metrics. Additionally, the correlations were higher for the alpha and beta bands. Finally, the correlations for the group-averaged connectomes were higher than the mean values of the correlations derived using individual-participant connectomes shown in Fig. 10.

**Figure 12:**
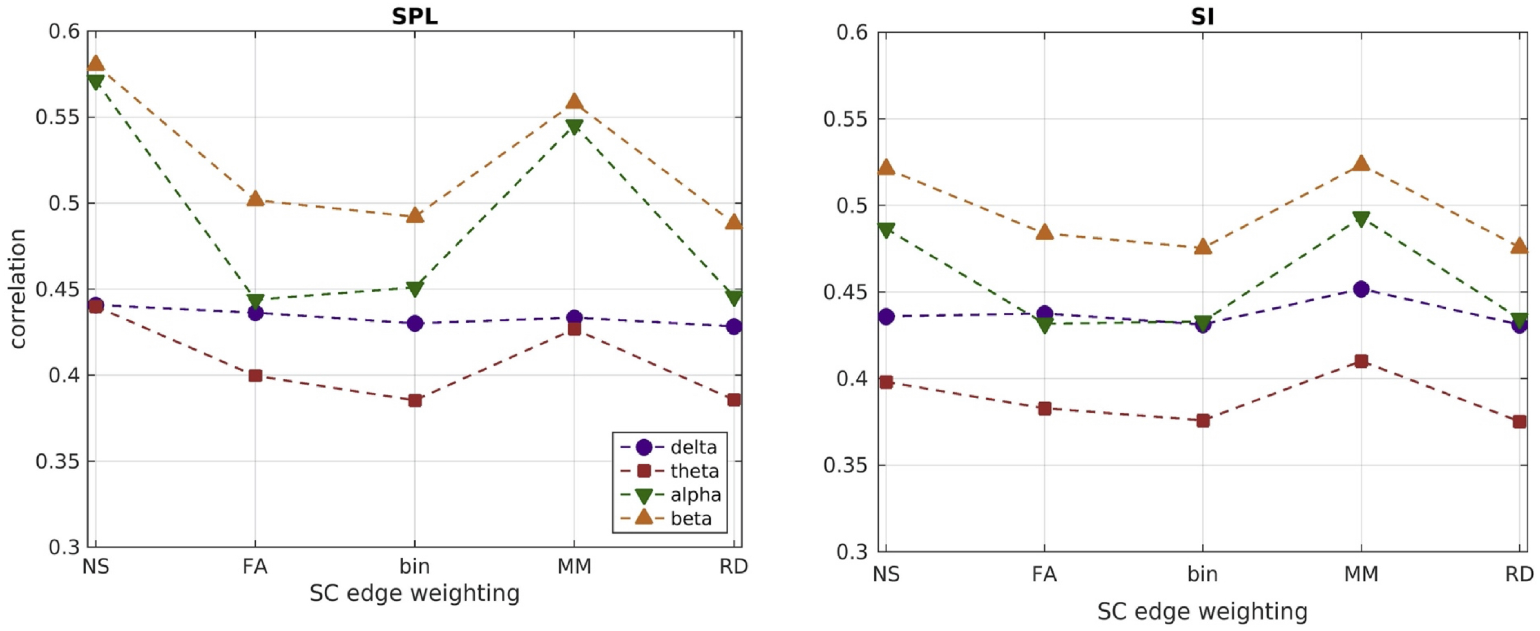
Correlations between predicted and observed FC, when the group-averaged structural and functional connectomes are used, for each of the two FC-predicting algorithms and the five structural edge-weightings, for each frequency band.

### 3.5 Shortest path length

The length of the shortest path (calculated from the weighted structural connectomes of each participant) for functionally connected brain areas depended on the metric used to weigh the edges of the SC matrices (Fig. 13). Weighting the SC matrices by NS or MM, which gave the highest correlations between FC_o_ and FC_p_, resulted in longer shortest-paths, compared to weighting them by FA or RD or using binarized graphs. With the NS- or MM-weightings, most shortest paths were 2 or 3 steps long, while for the other 3 edge-weightings they were 1 or 2 steps long.

**Figure 13:**
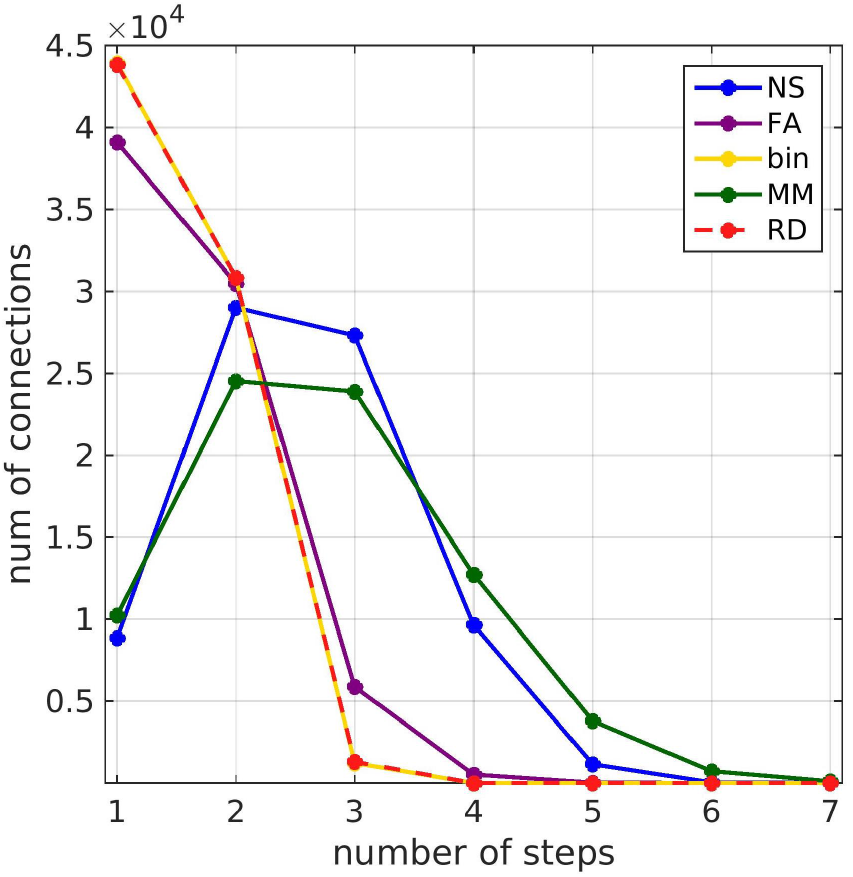
Histogram of the length of the shortest path between functionally connected brain areas, expressed as number of steps calculated based on the structural connectome, for the 5 different structural edge-weightings.

## 4 Discussion

Our work combines microstructural and electrophysiological brain imaging data to understand the relationship between brain structure and function in the human brain.

### 4.1 Novel contributions

The novel contributions of the work are as follows:

1. For each participant, we derived structural brain networks for which the edges are weighted with different attributes of the WM tracts that could influence functional connectivity, and binarized networks that only encode whether a structural connection exists or not. We used those structural networks in the search-information and the shortest-path-length algorithms (Gõni et al., 2014) to predict patterns of MEG resting-state functional connectivity, and we compared the resulting patterns. To the best of our knowledge, this is the first time that attributes of WM tracts beyond the number of streamlines and the mean fractional anisotropy are used in such algorithms, and the resulting patterns are compared. It is also the first time that these algorithms are used to predict MEG resting-state functional connectivity, and the first time that analysis aiming to describe the relationship between electrophysiological functional connectivity and structural connectivity is performed using individual-participant instead of averaged connectomes.
2. We applied, for the first time, NMF to the whole-scan MEG resting-state FC_o_ of our participants. This methodology uses the variance across participants to derive static connectivity components that comprise the cohort FC_o_.
3. We used the SI and SPL algorithms of Gõni et al. (2014) to calculate the FC_p_ both for the total FC_o_ and for each of its NMF components. We also calculated the correlations between the FC_o_ and the FC_p_, for the total observed functional connectivity and for each of its NMF components. This was done for each of the 5 different edge-weightings in the SC matrices. It reflects the reliability of the function predicting algorithms, and allows us to assess how accurate the different edge-weightings and function-prediction algorithms are at predicting patterns of FC that form the fundamental parts of the whole-scan resting-state MEG FC_o_ for this cohort. It also allows us to quantify the impact of the choice of structural edge-weighting and algorithm on the accuracy of the predictions when measuring electrophysiological functional connectivity by amplitude envelope correlations.
4. Motivated by the fact that the SPL algorithm was the best at predicting patterns of MEG resting-state functional connectivity in the majority of cases, we used the dMRI-derived structural connectome of each participant to calculate the shortest paths for each connection of the FC_o_, for all 5 structural egde-weightings.

### 4.2 Implications

The FC patterns predicted for a given participant, using different structural edge-weightings and algorithms, are similar to each other (Fig. 9), but there are exceptions where those patterns are not well correlated. This indicates that the choice of edge-weighting and algorithm has an impact on the FC_p_ and consequently on conclusions drawn from such studies. Our analysis also provides a quantification of the impact of the discrepancies that arise when calculating the FC_p_ with different structural edge-weightings, by calculating the correlations between FC_p_ and FC_o_. The distributions of correlations between FC_p_ and FC_o_ were statistically significantly different, and there is, for most NMF components, a significant improvement when either the NS or the MM is used to weight the edges of the structural networks. We also showed that for a given structural edge-weighting, the SPL algorithm resulted in higher correlation between the components of FC_p_ and the FC_o_ compared to the SI algorithm. This conclusion diverges from the observations of Gõni et al. (2014), who found that the SI algorithm could yield higher correlations. This could be due to the fact that they used fMRI rather than MEG data, as well as to the fact that they used averaged connectomes rather than participant-specific ones.

In order to put our work into context with respect to the existing literature we look at other studies that have compared structural networks to MEG-derived resting-state functional networks. Garcés et al. (2016) reported correlations in the range of 0.14 — 0.45 between group-averaged SC matrices and MEG-derived group-averaged FC matrices in the theta, alpha and beta bands. When considering individual-participant SC and FC matrices, the mean of the correlations was in the range of 0.04 — 0.25. These correlations are smaller than the correlations in our analysis, which were in the range of 0.35 — 0.60 for group-averaged connectomes and 0.20 — 0.31 for individual-participant connectomes. The differences are most likely due to the fact that Garcés et al. (2016) did not use any function-predicting algorithm, but rather compared structural and functional connectomes directly. Tewarie et al. (2014) reported adjusted R^2^ values of up to 0.18 for the linear fit of empirical MEG connectomes derived in the alpha band to a neural mass model applied on a binarized ‘backbone’ structural connectome derived in earlier work by Gong et al. (2009). This value of R^2^ is comparable to our correlation values of 0.5 — 0.55 for the two function-predicting algorithms in the alpha band (Fig. 12). The differences can be attributed to the use of the neural mass model as well as the fact that the structural connectome used by Tewarie et al. (2014) comes from a data set different to the one for which the MEG data were collected. Tewarie et al. (2019) reported R^2^ values between 0.25 — 0.46 when using the eigenmodes of the group-averaged structural connectome to predict group-averaged FC in the frequency bands we considered in our work. These values are, again, comparable to the correlation values we observed. Finally, Cabral et al. (2014) observed correlations in the range of 0.1 — 0.4 when using a coupled oscillator model to predict group-averaged MEG resting-state FC from structural connectivity. These correlations are lower than what we observe, likely due to the fact that the MEG data were collected from a cohort different to that from which the MRI data were collected. In general, using average connectomes could be valuable in that it allows for the most reproducible parts of the structural connectome to be used, and possibly discards some of the noise. At the same time it masks individual differences and therefore can a) artificially augment the correlations between the observed functional connectivity and that predicted by the analysis, as demonstrated in our work and, b) is not appropriate for the development of biomarkers that can allow classification of diseases. We also note that the study of (Tewarie et al., 2019) is in a sense reciprocal to ours: they decompose the structural connectome and use the derived basis to predict the observed functional connectivity, while we decompose the functional connectome and predict its components from the full structural connectome.

Brain areas that are close to each other and which are directly linked with a WM tract appeared to have stronger FC_o_. However, further analysis of the relationship between the distance of brain areas and whether or not they are linked or not revealed that it is the Euclidean distance that is the dominant predictor. The dependence of the strength of functional connectivity on whether brain areas are directly linked or not appears to be a result of the fact that areas that are closer together are more likely to have a WM tract directly connecting them. This would happen regardless of which brain atlas is used for parcellation.

Decomposing the FC_o_ using NMF revealed the components that form the complex pattern of the whole-scan FC_o_.

a. Delta band: The FC_o_ decomposed into components that have strong frontal connections and components that have strong occipital and parietal connections. There is also a separation of left-dominated from right-dominated activation. There is little variation in the mean of the correlations for different structural metrics and algorithms, for each of the 8 components of the FC_o_, and even less for the total FC_o_. All FC_o_ components exhibit higher correlations with the FC_p_ than the total FC_o_ exhibits with the FC_p_.
b. Theta band: The FC_o_ decomposed into components that are dominated by frontal and parietal inter-hemispheric connections, as well as fronto-parietal connections. For several of the components of FC_o_ the mean value of the correlations between the predicted and observed FC depends strongly on the edge-weighting used. The distributions over participants were statistically significantly different (corrected for multiple comparisons). For components 1, 3 and 6, the NS-weighted SC resulted in the highest correlations, while for components 2, 5, 7 and 8 the MM-weighting performed the best. For components 9 and 10, the FA resulted in a slightly higher mean value, however the differences with the mean values that resulted from the NS and MM weightings were not statistically significantly different (Table 12).
c. Alpha band: The FC_o_ decomposed into components that have predominantly inter-hemispheric occipital and parietal connections, as well as fronto-parietal connections (component 3) and temporo-occipital connections (component 6). The mean value of the correlations between the components of FC_o_ and FC_p_ was strongly dependent on the edge-weighting used. The corresponding distributions were statistically significantly different for all but a few of the pairs of SC metrics (Table 13 corrected for multiple comparisons via FDR). The MM and NS were the best predictors of FC_o_ for most of the components. The FA consistently performed poorly compared to those, and also compared to the binarized SC.
d. Beta band: The FC_o_ decomposed into components that comprise of mainly inter-hemispheric connections between occipital or parietal areas. There were also intra-hemispheric frontoparietal and temporo-parietal connections. There was a separation of left- and right-dominated connectivity in the 7^th^ and 8^th^ component. The mean value of the correlations between the components of FC_o_ and FC_p_ was strongly dependent on the edge-weighting used, with the corresponding distributions being statistically significantly different (corrected for multiple comparisons) for most the pairs of SC metrics (Table 14). The MM and NS were, again, the best predictors of FC_o_ for most of the components, while the FA consistently performed poorly.

Overall, the NS and MM edge-weightings were better predictors of the FC_o_ than the FA, which is frequently used in structural network analyses. In fact, the binarized SC matrices performed better than the FA-weighted ones in many cases. This finding, along with the fact that the FA has been shown to exhibit poor repeatability as an edge-weighting in other datasets (Messaritaki et al. (2019b),Messaritaki et al. (2019a)) suggest that the FA is of limited relevance when quantifying structural connectivity and trying to relate it to functional connectivity.

The correlations between the FC_p_ and the FC_o_, both for their non-decomposed values and for the components, were highly variable. For most of the components they were higher than the correlations that the total FC_o_ exhibited with the FC_p_. This indicates that the simplicity of the function-predicting algorithms that we considered is very good at capturing the dynamics of the fundamental components of the FC_o_, but perhaps less appropriate for understanding the whole-scan FC_o_. The correlations we observed for most of the NMF components are comparable, if slightly lower, to the correlations that were reported by Gõni et al. (2014), when they compared structural networks to fMRI-derived functional networks, even though they used average connectomes for the participants in their study. The correlations we observed for the whole-scan (pre-NMF) FC_o_ using the group-averaged connectomes were very similar to those of Gõni et al. (2014), which is a more approporiate comparison given their use of group-averaged connectomes.

Although our analysis used resting-state data, the documented correspondence between resting-state networks and task-related networks (Smith et al., 2009) indicates that the fundamental organization of the human brain is relatively similar across task and resting-state conditions. The validity of our conclusions should be explored for task-related studies too.

### 4.3 Assessment of our analysis

Head movement in the MEG system can have a detrimental effect on the measurement of restingstate brain activity, at both beam-former and network level, (Stolk et al., 2013; Messaritaki et al., 2017). To eliminate this confound, the method described by Stolk et al. (2013) and by Messaritaki et al. (2017) would have to be implemented. For that to be possible, continuous head localization throughout the MEG scan is necessary, so that the location of the fiducial points is recorded with the same frequency as the neuronal signals are recorded. However, this information was not recorded during the acquisition of the data. Despite that, the results of Messaritaki et al. (2017) show that the effect of head movement on MEG resting-state functional networks derived via an independent component analysis is small, indicating that the results are robust for head movement that is relatively low.

In our analysis, we considered one measure of MEG-derived functional connectivity, namely correlations between the Hilbert envelopes of the MEG beamformer in different brain areas. Our choice was motivated by the robustness of that measure (Colclough et al., 2016). Network behavior can also be characterised using phase-amplitude and phase-phase correlations of beamformer time series (Siegel et al., 2011), and those should be explored in similar analyses in the future.

Axonal diameter is an important factor when it comes to the formation of brain dynamics (Drakesmith et al., 2019). Including a direct measure of the distributions of axonal diameters in different WM tracts would have been desirable. However, this requires ultra-strong gradients (Veraart et al., 2020) and the data to be acquired with different sequences than those used here. For that reason we used the RD as a possible SC edge-weighting, because it has been shown to correlate with axon diameter (Barazany et al., 2009). Including such a measure in future analyses could result in more components of the FC_o_ having higher correlations with the FC_p_, as other mechanisms that promote FC are ultimately included in the models.

We only applied a very modest thresholding to the SC matrices. The thresholding of such matrices is an issue of debate nowadays. For example, Buchanan et al. (2020) showed that the spurious WM connections reconstructed with probabilistic tractography can be reduced by different thresholding schemes and give better age associations than if the connections were left unthresholded. On the other hand, Civier et al. (2019) evaluated the removal of weak connections from structural connectomes and concluded that the removal of those connections is inconsequential for graph theoretical analysis, ultimately advocating against the removal of such connections. Finally, Drakesmith, Caeyenberghs, Dutt, Lewis, David and Jones (2015) showed that thresholding can reduce the effects of false positives, but it can introduce its own biases. In our analysis, we used deterministic tractography, which results in fewer false positives (Sarwar et al., 2018). This, along with the fact that the function predicting algorithms we used favor stronger connections, indicate that any further thresholding is unlikely to have a significant impact on the patterns of FC_p_.

## 5 Conclusions

In this study, we used MRI and MEG data to investigate the relationship between brain structure and resting-state functional connectivity in the human brain. Function-predicting algorithms were shown to be good at predicting the building blocks of electrophysiological resting-state functional connectivity, as measured by MEG in our experiment. Different structural edge-weightings resulted in very different pictures for the structural connectome and for its relationship to electrophysiological functional connectivity. The number of streamlines and the myelin measure were the best predictors of the NMF components of the observed functional connectivity, when used with the shortest-path-length algorithm. Our results indicate that the accuracy of the predictions of similar studies can be improved if well-considered structural metrics are used in the analyses.

## 6 Acknowledgments

EM was partly funded by the BRAIN Biomedical Research Unit (which is funded by the Welsh Government through Health and Care Research Wales). EM is also funded by a Wellcome Trust ISSF3 Research Fellowship at Cardiff University (204824/Z/16/Z). SF was funded by CUBRIC and the School of Psychology, Cardiff University, and by an ISSF grant to Cardiff University School of Medicine. SS received funding from the University of Verona Internationalisation Programme 2019 (Action 4C). DKJ is funded by a Wellcome Trust Investigator Award (096646/Z/11/Z). DKJ and KDS were supported by a Wellcome Trust Strategic Award (104943/Z/14/Z). The work was also supported by a Wellcome Trust Strategic Award (100202/Z/12/Z), the MRC UK MEG Partnership Grant (MR/K005464/1) and the MRC Doctoral Training Grant (MR/K501086/1). LM and BR were funded by a MRC grant (MR/K005464/1). BR was also funded by MRC grant MR/K501086/1.

## 7 Competing Interests

The authors have no competing interests to disclose.

## Notes

### Competing Interest Statement

The authors have declared no competing interest.

